# Neuropeptide Y neurons of the locus coeruleus inhibit noradrenergic system activity to reduce anxiety

**DOI:** 10.1101/2023.10.16.562534

**Authors:** Danai Riga, Kelly Rademakers, Inge G. Wolterink-Donselaar, Frank J. Meye

**Affiliations:** Department of Translational Neuroscience, Brain Center, UMC Utrecht, Utrecht University, Utrecht, The Netherlands

**Keywords:** locus coeruleus, neuropeptide Y, noradrenergic system, neuronal excitability, neuromodulation, anxiolysis

## Abstract

Adaptive responses to challenging environments depend on optimal function of the locus coeruleus (LC), the brain’s main source of noradrenaline and primary mediator of the initial stress response. Built-in systems that exert regulatory control over the LC are largely unidentified. A good candidate system is neuropeptide Y (NPY), which is traditionally linked to anxiety-relief. Currently, the endogenous source of NPY to the LC, and how NPY-expressing neurons modulate the noradrenergic system to regulate anxiety remain unclear. We here identify, in mice, a novel NPY-expressing neuronal population (peri-LC_NPY_) neighboring LC noradrenergic neurons that locally innervates the pericoerulean space. Moreover, we demonstrate that stress engages peri-LC_NPY_ neurons, increasing their excitability. Mimicking peri-LC_NPY_ neuronal activation using *ex vivo* chemogenetics suppresses LC noradrenergic neuron activity, via an NPY Y1 receptor-mediated mechanism. Furthermore, *in vivo* chemogenetic stimulation of peri-LC_NPY_ neurons results in Y1R-dependent anxiety-relief. Conversely, inhibiting peri-LC_NPY_ neurons increases anxiety-like behaviors. Together, we establish a causal role for peri-LC_NPY_-mediated neuromodulation of the LC in the regulation of anxiety, providing novel insights in the endogenous mechanisms underlying adaptive responses to adversity.

## Introduction

Adaptive responses to stressful experiences are paramount to our survival and well-being. Opposing systems that mediate initiation and termination of the stress response work in tandem to ensure optimal adaptation to challenging environments^1,2^. Currently, knowledge of the factors that dictate adaptive modulation of the stress response is incomplete, limiting available therapeutics against stress-related afflictions, such as anxiety disorders^3,4^.

The locus coeruleus (LC) is the brain’s primary source of noradrenaline/norepinephrine (NE), and key regulator of the initial stress response^5–7^. The LC_NE_ system mediates arousal and allocation of attention, preparing the organism for task-relevant and salience-specific responses, required in novel environments commonly associated with high cognitive and emotional load^8–10^. To accomplish this, owing to its vast projection network and efferent collaterals^11^, the LC coordinates myriad functions, from sympathetic responses, such as heart rate and pupil dilation, to complex, high-order cognitive processes, including goal orientation and decision-making^9^. LC hyperactivity, characterized by high tonic firing, can lead to maladaptive responses to perceived threats, priming the development of pathological anxiety^7^. In support, optogenetic stimulation of LC_NE_ neurons results in anxiety in mice, whereas chemogenetic inhibition of LC_NE_ cells strongly suppresses stress-driven anxiety-like behaviors^12^. Given the importance of LC_NE_ activity in shaping anxiety responses, it is important to understand the mechanisms that appropriately regulate its function.

A strong candidate regulator of LC_NE_ activity is the neuropeptide Y (NPY) system, composed of groups of neurons that communicate via NPY release, and others that interpret these signals. NPY, one of the most widely distributed neuropeptides in the central nervous system, is traditionally associated with stress-coping and anxiety relief^13–15^ and it is dubbed the “stress resilience” molecule^3^ after early preclinical studies, which employed NPY or NPY receptor agonists, highlighted its anxiolytic effects^16–19^. In support, *NPY* gene expression as well as NPY plasma levels have been linked to trait anxiety and stress-related neuropsychological conditions in clinical settings^3,20^.

NPY immunoreactivity and NPY receptor presence have been observed in the LC^21,22^. However, few studies have examined the functional relationship between (endogenous) NPY-mediated neuromodulation and the LC_NE_ system. *Ex vivo* electrophysiological evidence shows that exogenously applied NPY reduces LC_NE_ spontaneous discharge^23^ and facilitates hyperpolarization of LC_NE_ neurons^24^. These studies highlight an inhibiting effect of NPY on tonic LC firing, which could be crucial in regulating LC activity during arousal, and under stress. In agreement, exogenous NPY application aiming at the peri-coerulean (peri-LC) space induces anxiolysis in the elevated plus maze (EPM)^25^, an innately anxiogenic behavioral task that is known to engage the LC^26^.

Despite these insights, the endogenous source of NPY to the LC remains unidentified. In the rat brain, NPY-like immunoreactivity has been observed in LC_NE_ cell bodies as well as projection fibers^21,27–30^, suggesting that NPY (co-)released by LC_NE_ neurons constitutes the main endogenous source of NPY to the LC. However, recent RNA sequencing data challenge this, demonstrating NPY presence in the peri-LC space but no co-expression in noradrenergic neurons of the mouse LC^31^. These contradictory reports have cast doubt on the origins of NPY input to the region. Furthermore, independently of its origins, no studies have addressed how endogenously released NPY modulates LC_NE_ activity to regulate anxiety levels. Understanding the effects of endogenous NPY signaling is critical, as pharmacologically applied NPY engages distinct NPY receptors, resulting in dose-dependent, opposing regulation of anxiety-like behaviors^32^.

Aiming to address this, we examined NPY organization and function in the mouse LC. We characterize a previously unidentified NPY-expressing, pericoerulean (peri-LC_NPY_) neuronal population at the anatomical, electrophysiological, and behavioral level. Our data indicate that peri-LC_NPY_ neurons constitute a distinct, non-noradrenergic population that is engaged by exposure to stress. We show that local peri-LC_NPY_ cells suppress LC_NE_ neuronal activity in an NPY-mediated, Y1 receptor-dependent manner. Finally, we demonstrate that activation of peri-LC_NPY_ cells reduces anxiety-like behaviors, via LC Y1 receptors, and conversely, peri-LC_NPY_ inhibition promotes anxiogenesis. Together, we here describe a previously unknown population of NPY-expressing cells that regulates the LC noradrenergic system, thereby promoting adaptive behavioral responses in arousing environments.

## Results

### Identification of NPY-expressing neurons in the pericoerulean space

To examine the presence of NPY neurons in the peri-LC region, *NPY-cre* mice^33^ were crossed with the *Ai14* reporter line^34^, enabling tdTomato (tdT) fluorescence selectively in NPY-expressing cells (Fig. S1A). We observed a clear presence of tdT^+^ cells neighboring the LC proper, the region occupied by noradrenergic cell bodies (Fig. 1A; Fig. S1B). We performed a systematic mapping of peri-LC_NPY_ neuron location with respect to noradrenergic neurons of the LC (LC_NE_), identified by tyrosine hydroxylase (TH) expression. In coronal slices immunolabelled for TH, we extracted the coordinates of tdT^+^ cells at the entire rostrocaudal axis (Anterior-Posterior: -5.80 to -5.25 mm from bregma) containing the LC (Fig. 1A). We then used the distance from LC center to plot peri-LC_NPY_ cell distribution at the mediolateral (Fig. 1B, Fig. S1C) and dorsoventral axes (Fig. 1C, Fig. S1C). Within the pericoerulean region, defined by the extend of reach of LC dendritic processes (Fig. S1B)^35^, peri-LC_NPY_ neurons congregated largely medially (∼60%) to the LC proper, with ∼20% found dorsal and ∼40% ventral to TH^+^ cell bodies.

**Figure 1.**
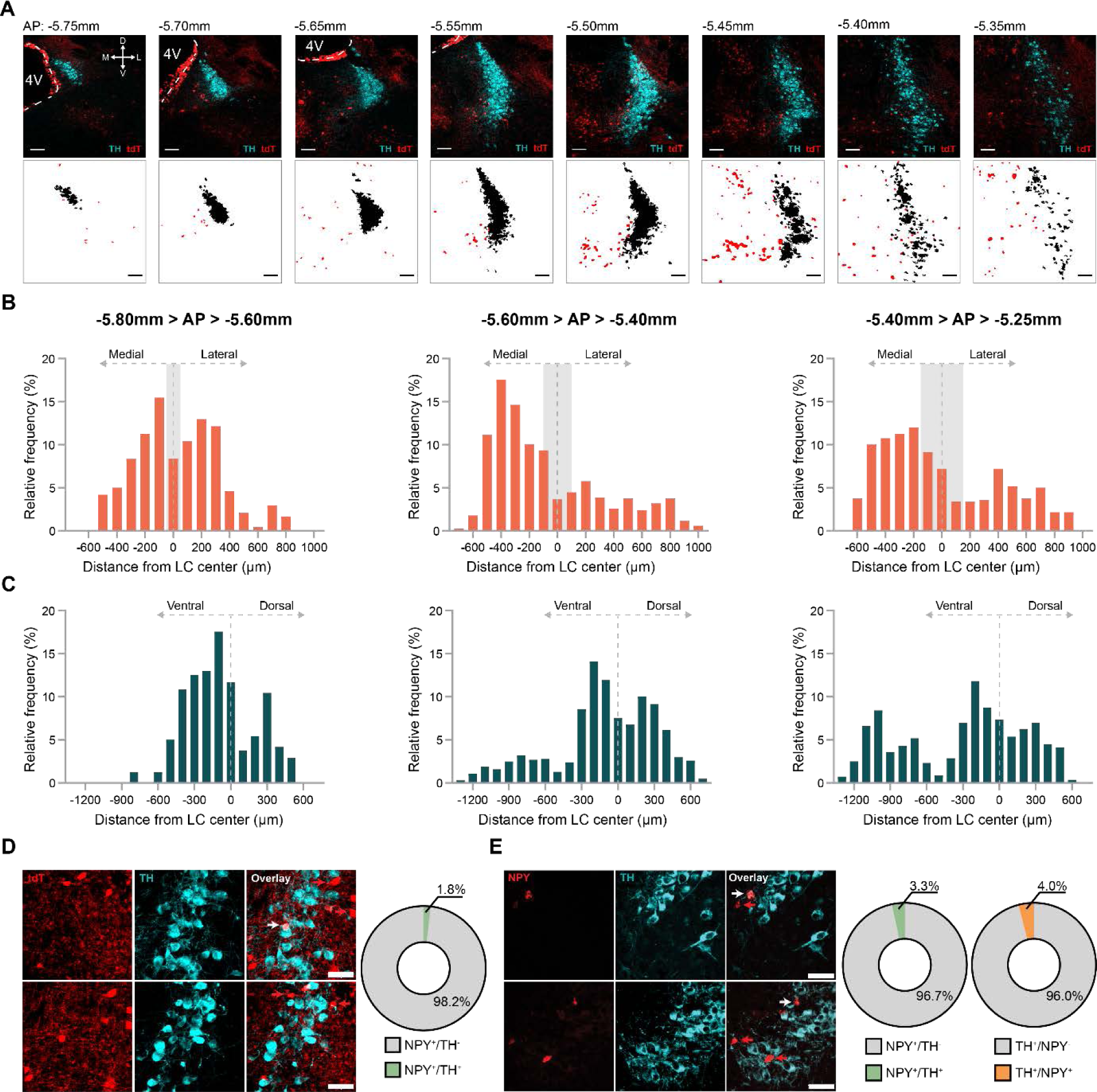
Peri-LC_NPY_ neurons are distributed medial to LC proper and are TH-lacking. A) Top: Representative examples of tdTomato expression (tdT, red) and noradrenergic immunolabeling (TH, cyan) in coronal slices from *NPY-cre:Ai14* mice, depicting the LC at the entire rostrocaudal axis. 4^th^ ventricle is indicated. Bottom: FIJI-processed images containing masks for the two channels (tdT, red; TH, black), on which detection and coordinate extraction per tdT^+^ cell was based. Scale bar, 100µm; D, dorsal; V, ventral; M, medial; L, lateral. B) Frequency of distribution (%) of peri-LC_NPY_ neurons location in respect to their distance (µm) from LC center at the mediolateral axis. Grey shading depicts the width of LC proper in corresponding coordinates. Across the AP axis, the majority of peri-LC_NPY_ cells, are located medially to LC proper. C) Frequency of distribution (%) of peri-LC_NPY_ neurons location in respect to their distance (µm) from LC center at the dorsoventral axis. Across the AP axis, the majority of peri-LC_NPY_ cells are located ventral to LC proper. D) Left: Representative examples of tdTomato expression (tdT, red) and noradrenergic immunolabeling (TH, cyan) in coronal slices from *NPY-cre:Ai14* mice. Right: Quantification of double-immunoreactive NPY^+^/TH^+^ cells showed that peri-LC_NPY_ neurons are predominantly TH-lacking. Red arrows, NPY^+^/TH^-^ cells (98.2%); white arrow, NPY^+^/TH^+^ cells (1.8%); Scale bar, 50µm. E) Left: Representative examples of *NPY* expression (red), detected by in situ hybridization, and noradrenergic immunolabeling (TH, cyan) in coronal slices from wild-type C57BL/6 mice, containing the LC. Right: Quantification of *NPY* puncta and TH expression showed that the majority of peri-LC_NPY_ neurons are non-noradrenergic (NPY^+^/TH^-^, 96.7%, red arrow; NPY^+^/TH^+^, 3.3%, white arrow), and vice-versa, only 4% of LC_NE_ cells co-express *NPY* (TH^+^/NPY^-^ 96%, red arrows; TH^+^/NPY^+^, 4%, white arrow). Scale bar, 50µm. B-C: AP, -5.80mm to -5.60mm, N=2 mice, n=239 cells; AP, -5.60mm to -5.40mm, N=7 mice, n=1004 cells; AP, -5.40mm to -5.25mm, N=6 mice, n=558 cells. D) N=5 mice, n=612 cells. E) *NPY*: N=2 mice, n=30 cells; TH: N=3 mice, n=125 cells.

Our mapping data indicated that, based on location alone, the peri-LC_NPY_ population is, to a large degree, topographically distinct from LC_NE_ cells. To further validate this, in coronal slices from *NPY-cre:Ai14* mice immunolabelled for TH, we quantified the percentage of TH-expressing peri-LC_NPY_ cells in the total peri-LC_NPY_ population (Fig. 1D). In contrast to earlier studies, which showed large overlap between noradrenergic and NPY-expressing neurons^21,27^, the vast majority of tdT^+^ cells were TH-devoid, while only 1.8% co-expressed TH, indicating that the absolute number of NPY neurons occupying the pericoerulean space has been largely underestimated^31^. To control for incomplete Cre-recombinase expression in *NPY-cre:Ai14* mice that could potentially confound these results, we used multiplex fluorescent RNAscope in situ hybridization against endogenous *NPY*, in combination with TH immunolabeling, in brain slices from wild-type C57BL/6 mice (Fig. 1E). We quantified TH expression in *NPY*^+^ cells and vice-versa, *NPY* puncta in LC_NE_ neurons. In agreement with our earlier results, only 3.3% of peri-LC_NPY_ neurons co-expressed TH, while 4% of LC_NE_ cells were NPY^+^.

### Peri-LC_NPY_ neurons respond to acute stress

The LC is an important regulator of the initial stress response, with implications for the development of stress-induced pathology^7^. To further understand potential contributions of peri-LC_NPY_ neurons in neuromodulation of the LC_NE_ system, we examined their physiological properties under naïve conditions and after stress. For this, we performed whole-cell patch clamp recordings in LC-containing brain slices of control or stressed *NPY-cre:Ai14* mice (Fig. 2D, Fig. S2A). First, we established a stress protocol that induces acute and long-lasting anxiety-like phenotypes, following exposure to electrical foot-shocks (Fig. 2A-C). Acutely (30 min) after stress, foot-shocked exposed mice (FS) showed decreased time spent in the open arm of an elevated-plus maze (EPM) compared to non-stressed (NS) controls (NS, 13.2% *vs.* FS, 6.9% of total exploration time, Fig. 2B). This was accompanied by an increase in anxiety index (NS, 0.76 *vs.* FS, 0.84), a compound parameter that integrates avoidance and exploratory behaviors^36^, further supporting direct stress effects on anxiety-like behaviors^6^. Stress-induced anxiety was long-lasting, as reflected in reduced exploration of an open field one week following exposure to foot-shocks (Fig. 2C). In particular, FS mice spent less time at the center of the open field arena (NS, 4.0% *vs.* FS, 2.4% of total exploration time), which they visited less frequently compared to controls (NS, 24.5 *vs.* FS, 17.5 visits).

**Figure 2.**
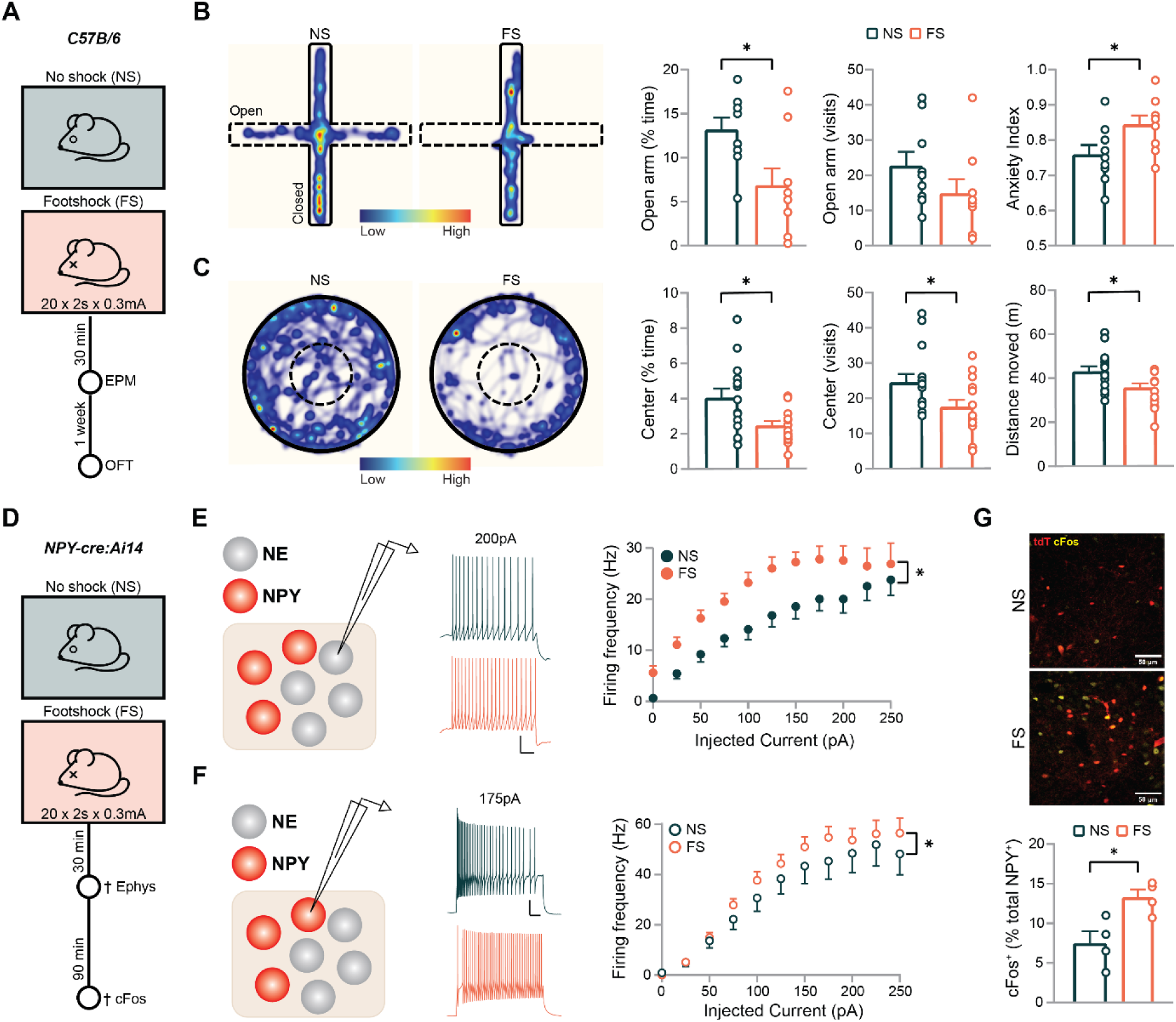
Peri-LC_NPY_ neurons are stress-responsive. A) Schematic representation of experimental design. Acute (30 min, elevated plus maze -EPM) and long-lasting (1-week, open field test -OFT) effects of stress (electrical foot-shock: 20 shocks x 2s x 0.3mA) were assessed in wild-type C57BL/6 mice. B) Stress exposure acutely increased anxiety-like phenotypes in the EPM. Stressed mice spent less time in the open arms of the maze when compared to controls (NS, N=9, 13.2% *vs.* FS, N=9, 6.9% of total EPM time; unpaired t-test, t(16)=2.694, *P*=0.016). Although decreased, no statistically significant group effect on the frequency of visits to the open arms was seen (NS, 22.7 *vs.* FS, 14.9; Mann-Whitney U=22, *P*=0.108). The stress group displayed increased anxiety index, which integrates explorative behavior in addition to avoidance in the assessment of anxiety-like behaviors^36^ (NS, 0.76 *vs.* FS, 0.84; unpaired t-test, t(16)=2.306, *P*=0.035). C) Stress exposure led to long-lasting anxiety-like phenotypes in the OFT. The FS group spent less time in the center of the open field arena (NS, N=15, 4.0% *vs.* FS, N=15, 2.4% of total OFT time; unpaired t-test, t(28)=2.727, *P*=0.011) and displayed reduced frequency of visits to the center (NS, 24.5 *vs.* FS, 17.5; unpaired t-test, t(28)=2.286, *P*=0.030). Stress had a lingering effect on explorative behavior in the OFT arena (Distance moved: NS, 43m *vs.* FS, 35.7m; unpaired t-test, t(28)=2.486, *P*=0.019). B-C: Representative spatial location heatmaps show time spent exploring the arenas. D) Schematic representation of experimental design. Control (NS) or foot-shock-subjected (FS) *NPY-cre:Ai14* mice were sacrificed 30 min following stress exposure for electrophysiological recordings. In a different cohort, NS and FS mice were sacrificed 90 min post-stress to examine cFos expression via immunolabeling. E) Whole-cell patch clamp recordings from LC_NE_ neurons. Stress exposure increased LC_NE_ firing frequency, as seen in the number of action potentials fired in response to increasing current injections (2-way RM ANOVA, main stress effect: F(1,26)=9.27, *P*=0.005). Representative example traces of action potentials fired in response to a 200pA current injection in NS (top) and FS (bottom) mice are depicted. F) Whole-cell patch clamp recordings from tTd^+^ peri-LC_NPY_ neurons were conducted in current-clamp configuration, to examine their intrinsic electrical properties, and possible effects of acute stress. Stress significantly increased peri-LC_NPY_ neuron firing frequency (Hz), observed in response to increasing current injections (2-way RM ANOVA main group effect, F(1,922)=5.597, *P*=0.018. Representative example traces of action potentials fired in response to a 175pA current injection in NS (top) and FS (bottom) mice are depicted. G) Quantification of cFos-expressing peri-LC_NPY_ cells showed increased percentage of double-immunoreactive neurons after exposure to stress (NS 7.5%, FS 13.3% of total peri-LC_NPY_ population; unpaired t-test, t(6)=3.169, *P*=0.019). Representative examples of tdTomato expression (tdT, red) and cFos immunolabeling (yellow) in coronal slices are shown. Scale bar, 50µm. B) NS, N=9; FS, N=9. C) NS, N=15; FS, N=15. E) NS, N=5 mice, n=12 cells, FS, N=6 mice, n=16 cells. F) NS, N=10 mice, n=40 cells; FS, N=13 mice, n=46 cells. G) NS, N=4 mice, n=1572 cells, FS, N=4 mice, n=867 cells. E, F) Scale bar: 20 mV, 100 ms. Data depicted as mean ± SEM. * *P* <0.05.

We verified that this stress protocol engages LC_NE_ cells. We performed patch-clamp recordings from LC_NE_ cells in brain slices from mice previously subjected to the foot shock paradigm or control conditions. LC_NE_ cells were identified based on their location and morphological characteristics^39^.

To validate that this approach yielded recordings from putative LC_NE_ neurons, we first confirmed the identity of a subset of biocytin-filled recorded cells using post-hoc TH immunolabeling (Fig. S2A). Our electrophysiological recordings demonstrated that foot shock experience engaged LC_NE_ cells, increasing their excitability. In particular, in slices prepared from stressed mice, we observed a higher number of action potentials in response to increasing current injections, as compared to controls (Fig. 2E). This was accompanied by a reduction in the current necessary for LC_NE_ cells to exceed their action potential threshold and fire (Rheobase: NS, 25pA, FS, 0pA, Fig. S2B), further corroborating increased neuronal excitability after stress.

We next recorded peri-LC_NPY_ intrinsic properties in slices prepared from the same cohort of control and stress-exposed mice (Fig. 2D, S2C). Stress did not affect peri-LC_NPY_ passive properties, such as cell capacitance and membrane resistance (Fig. S2D). Likewise, action potential threshold and resting membrane potential remained unaltered after exposure to stress (Fig. S2D). Control peri-LC_NPY_ neurons showed sustained capacity for high-frequency firing rates (Fig. 2F), corresponding well to the physiological requirements for neuropeptidergic release from dense core vesicles^37,38^. Notably, in brain slices prepared from stressed mice we detected an even greater number of action potentials in response to increasing current injections, as compared to controls (Fig. 2F), indicating peri-LC_NPY_ neuron engagement in stressful environments. To further validate this, we quantified cFos expression, as proxy for neuronal activation, in peri-LC_NPY_ cells from control and stress-exposed mice (Fig. 2G, Fig. S2E). We found an increased number of cFos^+^ peri-LC_NPY_ cells in mice subjected to foot-shocks (NS, 7.5%; FS 13.3% of total peri-LC_NPY_ population), corroborating the involvement of peri-LC_NPY_ neurons in the initial stress response.

### Peri-LC_NPY_ neurons innervate the pericoerulean space but do not form GABAergic or glutamatergic synaptic connections with LC_NE_ neurons

We demonstrated that exposure to an acute stressor simultaneously engages LC_NE_ and peri-LC_NPY_ neurons, indicating an involvement of the two populations in the stress response. However, the effects of peri-LC_NPY_ activation on the LC_NE_ system remained unknown. To examine whether peri-LC_NPY_ neurons could provide (NPY) input to the LC proper, we employed Cre-dependent, virus-mediated antero-and retrograde labeling of peri-LC_NPY_ cells and mapped their neuroanatomical circuitry^40^. The vast majority of NE dendritic processes are located outside the nuclear core, where they receive extensive non-noradrenergic synaptic contacts^35^. Thus, we first investigated whether peri-LC_NPY_ neurons project within the dendritic zone of the LC, to permit signaling from NPY and its co-transmitters in the region.

For this, *NPY-cre* mice were bilaterally injected with a Cre-dependent AVV (AAV-Syn-FLEX-CoChR-GFP) in the LC and allowed a period of ≥5 weeks of virus incubation. Next, brain slices were collected and a qualitative analysis of labeled NPY^+^ cell bodies and fibers locally, within the LC, and in selected projection fields, was performed (Fig. 3A, Fig. S3A). While we observed clear innervation of the pericoerulean space, we detected no peri-LC_NPY_ efferents in other evaluated brain regions with known LC outputs (Fig. 3A, Fig. S3A). This suggests that local peri-LC_NPY_ neurons do not possess long-range projection neuron characteristics, as seen for instance in cortical areas^41^. To further corroborate local connectivity of peri-LC_NPY_ neurons within the region, we next injected a Cre-dependent retrograde HSV virus (HSV-hEF1a-LS1L-mCherry) in the LC. This resulted in retrograde tracing of NPY^+^ cell bodies throughout the pericoerulean area (Fig. 3B, Fig. S3B), supporting the notion that peri-LC_NPY_ neurons terminate within the region. Moreover, we detected several brain areas containing LC-projecting NPY^+^ cells, expanding previously reported data on NPY-containing afferents^42^ (Fig. S3B, Table S1). Both viral tracing strategies indicated that peri-LC_NPY_ projections assemble in the pericoerulean region. To assess whether these fibers are terminating or fibers of passage, we bilaterally injected an AAV construct (AAV-hSyn1-mCBP-EGFP-2A-mSyp1-mRuby) in the LC of *NPY-cre* mice for Cre-dependent expression of membrane-bound GFP and mRuby-fused synaptophysin, that enables axonal and presynaptic terminal labeling, respectively^43^ (Fig. 3C). Within the LC dendritic zone, we identified mRuby^+^ puncta along peri-LC_NPY_ axons, suggesting LC_NE_ synaptic innervation by local peri-LC_NPY_ cells.

**Figure 3.**
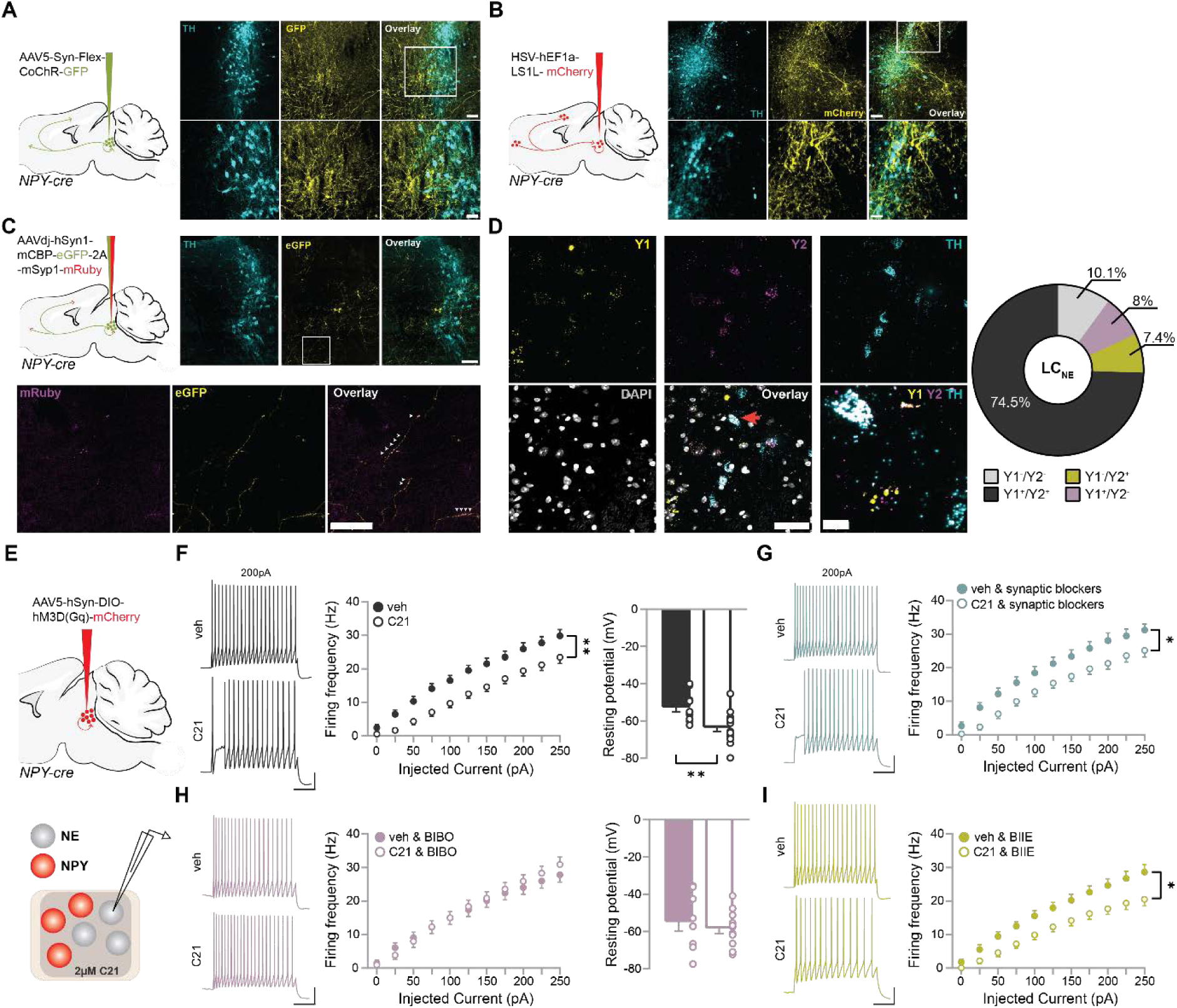
Peri-LC_NPY_ neurons reduce LC_NE_ excitability via Y1Rs. A) Left: Schematic of sagittal mouse brain, depicting the location of bilateral virus injections for anterograde labeling in *NPY-cre* mice. Right: Representative images of GFP expression (yellow) and noradrenergic (TH, cyan) immunolabeling in coronal slices containing the LC, illustrating dense NPY innervation in the peri-LC space. B) Left: Schematic of sagittal mouse brain, depicting the location of bilateral virus injections for retrograde labeling in *NPY-cre* mice. Right: Representative images of mCherry expression (yellow) and noradrenergic (TH, cyan) immunolabeling in coronal slices containing the LC, illustrating retrogradely-traced NPY^+^ cell bodies and fibers in the peri-LC space. C) Left: Schematic of sagittal mouse brain, depicting the location of bilateral virus injections for axonal and presynaptic labeling in *NPY-cre* mice (N=5). Right: Representative images of GFP expression (yellow) and noradrenergic (TH, cyan) immunolabeling in coronal slices containing the LC, illustrating NPY^+^ cell bodies and axons in the pericoerulean space. Inset: mRuby^+^ puncta are traced along NPY^+^ axons (white arrow heads), indicating that peri-LC_NPY_ neurons synapse within the LC dendritic zone. D) Left: Representative examples of Y1R (yellow), Y2R (magenta), and TH (cyan) mRNA detected by in situ hybridization, in coronal slices from wild-type C57BL/6 mice containing the LC. Right: Quantification of Y1R and Y2R puncta showed that the majority of LC_NE_ neurons expressed both NPY receptors (Y1^+^/Y2^+^, 74.5% of detected cells, orange arrow and expanded panel). 8% of LC_NE_ neurons contained Y1Rs, 7.4% Y2Rs and 10.1% were YR-lacking. E) Schematic of sagittal mouse brain, depicting the location of bilateral virus injections used to drive hM3Dq expression in *NPY-cre* mice. After a period of virus incubation (≥ 5 weeks), brain slices were prepared for electrophysiological recordings. Whole-cell patch clamp recordings from LC_NE_ neurons were conducted in current-clamp configuration, in presence of the DREADD actuator C21 (2µM) or vehicle. F) In presence of C21 we detected fewer action potentials in response to increasing current injections, as compared to vehicle (2-way RM ANOVA, main effect of treatment, F(1,30)=10.36, *P*=0.003). Chemogenetic activation of peri-LC_NPY_ neurons induced membrane hyperpolarization (resting membrane potential: veh, -52.7mV; C21, -63.5mV; unpaired t-test, t(23)=3.30, *P*=0.003). G) Similar effects of C21 bath application were observed in presence of synaptic blockers against AMPAR (10µM CNQX), NMDAR (50µM D-AP5), GABA_A_R (100µM picrotoxin) and GABA_B_R (10µM CGP-54626) -mediated currents (2-way RM ANOVA, main effect of treatment, F(1,25)=6.18, *P*=0.020). H) Pretreatment with the selective Y1R antagonist BIBO-3304 (1µM) blocked the effect of peri-LC_NPY_ chemogenetic activation on LC_NE_ firing frequency (2-way RM ANOVA, main effect of treatment, F(1,24)=0.06, *P*=0.815) and membrane hyperpolarization (resting membrane potential: veh, -55.0mV; C21, -60.6mV; unpaired t-test, t(16)=1.74, *P*=0.100). I) Pretreatment with the selective Y2R antagonist BIIE-0246 (1µM) did not preclude the effects of C21 on LC_NE_ excitability (2- way RM ANOVA, main effect of treatment, F(1,21)=7.41, *P*=0.013). A-C: Scale bar, 100µm and 50µm (inset). D) Scale bar, 50µm and 10µm (expanded panel). F-I) Representative traces of LC_NE_ firing in response to 200pA current injection (scale bar 20mV, 200 ms). A) N=5. B) N=5. C) N=3. D) N=4 mice, n=149 cells. F) Veh, N=4 mice, n=15 cells; C21, N=4 mice, n=17 cells. G) Veh & block, N=2 mice, n=14 cells; C21 & block, N=3 mice, n=13 cells. H) Veh & BIBO-3304, N=3 mice, n=13 cells; C21 & BIBO- 3304, N=4 mice, n=13 cells. I) Veh & BIIE-0246, N=2 mice, n=11 cells; C21 & BIIE-0246, N=3 mice, n=12 cells. F-I). Data depicted as mean ± SEM. * *P* <0.05; ** *P* <0.01.

Based on these data, we hypothesized that peri-LC_NPY_ neurons form functional synapses with LC_NE_ cells, necessary for LC_NE_ neuromodulatory control. To test this, we bilaterally injected *NPY-cre* mice with a Cre-dependent AAV (AAV-Syn-FLEX-CoChR-GFP) in the LC, that enables selective expression of the highly-conducting channelrhodopsin variant CoChR^44^ in peri-LC_NPY_ neurons (Fig. S4A, B). After a period allowing for virus incubation and CoChR expression in terminals (≥5 weeks), we performed whole-cell patch clamp recordings from putative LC_NE_ neurons (*cf*., Fig. S2A) To investigate the presence of direct peri-LC_NPY_◊LC_NE_ synaptic connectivity we opto-stimulated peri-LC_NPY_ neurons and recorded from LC_NE_ cells in brain slices with confirmed CoChR innervation (Fig. S4C, D).

First, to detect AMPAR-mediated or GABA_A_R-mediated ionotropic currents, we optogenetically stimulated peri-LC_NPY_ neurons with single pulses, while recording from LC_NE_ cells in voltage clamp configuration at -50mV. The fraction of LC_NE_ neurons that displayed perceptible postsynaptic responses to peri-LC_NPY_ optostimulation was negligible (AMPA-mediated, 0/39 cells; GABA_A_R-mediated, 1/39 cells). Rather, the majority of LC_NE_ neurons remained unresponsive (38/39 cells, Fig. S4C). In these experiments we also applied trains of 20 pulses of 5, 20 or 50 Hz photostimulation, to allow for the possibility of high frequency stimulation requirements in detecting forms of (GABA_B_R or NPY receptor-mediated) metabotropic signaling^45,46^. As above, no such direct fast onset responsivity was observed between peri-LC_NPY_ and LC_NE_ neurons (Fig. S4C).

Because we performed the experiments above with a potassium gluconate-based intracellular solution, we may have underestimated connectivity at more distal inputs^47^ to LC_NE_ neurons. To address this, we next recorded a subset of LC_NE_ cells using a cesium chloride (CsCl)-based internal in the absence of synaptic blockers, at -60mV, to allow for the detection of GABA_A_R or eventual AMPAR-mediated currents^47^. In agreement with our previous data, no responses to single optical pulse were observed (20/20 cells, non-responsive, Fig. S4D). Finally, to account for the possibility of silent synapses, we performed recordings at +40mV using the CsCl-based internal solution to detect NMDAR-mediated currents. As before, no synaptic responses to peri-LC_NPY_ stimulation were seen (12/12 cells, non-responsive, Fig. S4D). Taken together, these experiments indicate that there is close to null glutamatergic or GABAergic synaptic connectivity (either ionotropic nor fast-onset metabotropic) between peri-LC_NPY_ and LC_NE_ cells.

### Pharmacologically applied NPY bidirectionally alters LC_NE_ neuronal excitability, via distinct NPY receptors

Our connectivity data suggest that any potential communication between peri-LC_NPY_ and LC_NE_ neurons is likely mediated by NPY itself. Although we cannot exclude that NPY signaling could manifest in our functional connectivity studies, to our knowledge fast-onset synaptically-driven, direct postsynaptic NPY receptor currents have not been observed before^47^. Thus, we reasoned that peri-LC_NPY_-LC_NE_ direct NPY signaling occurs in an alternative manner, the detection of which cannot be achieved by optogenetic circuit mapping approaches. To further investigate this possibility, we first examined NPY receptor presence in LC_NE_ cells, which would be required to mediate direct NPY signaling. Using multiplex fluorescent RNAscope in situ hybridization we identified LC_NE_ neurons that expressed Y1R and/or Y2R puncta in wild-type C57BL/6 mice (Fig. 3D). In accordance with previous reports^30^, the majority of LC_NE_ neurons expressed both receptors (Y1^+^/Y2^+^, 74.5%). Y1R (Y1^+^/Y2^-^, 8%) or Y2Rs (Y1^-^/Y2^+^, 7.4%) -expressing LC_NE_ subgroups were observed at a lesser extent, while the remaining fraction was YRs-lacking (Y1^-^/Y2^-^, 10.1%). Together, our data verified that LC_NE_ neurons express the necessary molecular machinery for NPY-mediated neuromodulation.

Next, we investigated whether YR-mediated NPY signaling alters LC_NE_ intrinsic excitability. For this, we performed whole-cell patch clamp recordings in the presence of different doses of NPY. In brain slices prepared from wild-type C57BL/6 mice, we bath-applied NPY or vehicle, and recorded LC_NE_ firing frequency. In the presence of 30nM NPY we detected fewer action potentials in response to increasing current injections, as compared to vehicle. This effect was mediated by Y1Rs, as NPY-induced decrease in LC_NE_ excitability was blocked in slices pretreated with the selective Y1R antagonist BIBO-3304^48^ (Fig. S5A). Surprisingly, a 10-fold increase in NPY concentration yielded the opposite result, as we observed LC_NE_ hyper-excitability in NPY-treated slices. Notably, LC_NE_ increased firing was abolished in the presence of the selective Y2R antagonist BIIE-0246^49^ (Fig. S5B). Furthermore, 300 nM NPY-induced LC_NE_ hyper-excitability was prevented in slices pre-treated with synaptic blockers (for AMPA/kainate, NMDA, GABA_A_ and GABA_B_ receptors), suggesting that changes in LC_NE_ excitability may occur via an indirect network effect (Fig S5C). Overall, these observations highlight the capacity of LC_NE_ neurons to respond directly and indirectly, via distinct receptors, to pharmacologically applied NPY, with opposing effects on their excitability. This emphasizes the need to understand the mechanisms underlying endogenous NPY-mediated neuromodulation of the LC_NE_ system.

### Peri-LC_NPY_ neurons suppress LC_NE_ excitability via postsynaptic Y1Rs

Our pharmacological data demonstrated that NPY has the capacity to bidirectionally alter LC_NE_ excitability states. However, whether endogenous, peri-LC_NPY_-mediated signaling could affect LC_NE_ firing properties remained unknown. To address this, we combined chemogenetic activation of peri-LC_NPY_ neurons, which allows for protracted peri-LC_NPY_ stimulation, with *ex-vivo* slice electrophysiology and assessed its effects on LC_NE_ firing patterns. To this end, *NPY-cre* mice were bilaterally injected with a Cre-dependent AAV (AAV-hSyn-DIO-hM3D(Gq)-mCherry) to drive the expression of a Gq-coupled (excitatory) designer receptor exclusively activated by designer drugs (DREADD) in peri-LC_NPY_ cells (Fig. 3E). This viral construct enabled targeted chemogenetic activation of peri-LC_NPY_ neurons in presence of the DREADD agonist compound 21 (C21)^50^. First, we validated that C21 activates peri-LC_NPY_ neurons *ex vivo* (Fig. S6A). Then, we proceeded with whole-cell patch clamp recordings of LC_NE_ neurons in brain slices prepared from *NPY-cre* mice bilaterally expressing the hM3Dq DREADD (Fig. 3E).

Passive electrophysiological properties and intrinsic excitability of LC_NE_ cells were assessed in current-clamp configuration, after bath application (≥10 min) of C21 (2µM) or vehicle. In the presence of C21 we detected fewer action potentials in response to increasing current injections as compared to vehicle (Fig. 3F), mimicking the effect of low NPY dose application. This effect was accompanied by LC_NE_ membrane hyperpolarization (Resting membrane potential: vehicle, - 52.7mV; C21, -63.5mV) and an increased rheobase (vehicle, 15.0pA; C21, 44.1pA; Fig. 3F, Fig. S6B), indicating that activation of local peri-LC_NPY_ neurons results in reduced LC_NE_ neuronal excitability. Peri-LC_NPY_ chemogenetic stimulation did not alter other electrophysiological parameters of LC_NE_ cells (Fig. S6B). Importantly, in brain slices that lacked hM3Dq expression in the region, we confirmed that C21 did not result in off-target, non-specific effects on LC_NE_ neuronal firing (Fig. S6C).

We next addressed the signaling mechanisms through which peri-LC_NPY_ neuron activity suppresses the excitability of LC_NE_ cells. Particularly, we sought to address (i) the likelihood of the peri-LC_NPY_ stimulation effect being direct or indirect via intermediate (e.g., GABAergic^51^) neurons; and (ii) whether it could be mediated by peri-LC_NPY_ neurons co-releasing signaling molecules other than NPY, such as GABA or glutamate, despite the latter seeming unlikely (*cf.,* Fig. S4C, D). To that end, we repeated the experiment of chemogenetic peri-LC_NPY_ neuron stimulation, while assessing LC_NE_ neuron excitability, but this time we pharmacologically blocked synaptic network activity (i.e., blocking AMPAR/ kainateR, NMDAR, GABA_A_R and GABA_B_Rs; *cf.,* Fig. S5C). In these conditions, we still observed a clear decrease in LC_NE_ firing in presence of C21 (Fig. 3G), pointing towards direct post-synaptic effects of peri-LC_NPY_ neuron stimulation on LC_NE_ cell excitability.

Finally, to examine whether C21 effects were specifically mediated by NPY signaling, we assessed LC_NE_ excitability in presence of Y1R or Y2R blockers. In slices pretreated with the Y1R antagonist BIBO-3304 (1µM), the effects of C21 on LC_NE_ excitability and membrane hyperpolarization were fully occluded (Resting membrane potential: vehicle, -55.0mV; C21, -60.6mV; Fig. 3H). On the contrary, blocking Y2Rs with BIIE-0246 (1µM) did not prevent the ability of peri-LC_NPY_ stimulation to reduce LC_NE_ firing frequency (Fig. 3I). Synaptic blockers, BIBO-3304 or BIIE-0246 alone did not alter LC_NE_ firing properties in vehicle-treated slices, further highlighting the specificity of the effects of peri-LC_NPY_ chemogenetic activation (Fig. S6D).

Together, our data suggest that sustained stimulation of local peri-LC_NPY_ neurons hyperpolarizes LC_NE_ cells and dampens their excitability. Our observations point to postsynaptic Y1Rs, situated onto LC_NE_ neurons, being the prime mediators of local NPY signaling on the noradrenergic system. Notably, C21-driven activation of peri-LC_NPY_ neurons and exogenous application of 30 nM NPY (but not of 300 nM NPY) were analogous in regard to their effects on LC_NE_ intrinsic excitability, offering insights regarding the potential concentrations of chemogenetically-evoked NPY release from peri-LC_NPY_ cells.

### Peri-LC_NPY_ neurons bidirectionally modulate anxiety-like behavior via local Y1Rs

Optogenetic LC_NE_ neuron stimulation is anxiogenic at baseline conditions, and conversely chemogenetic LC_NE_ neuron silencing prevents stress-induced anxiety^12,52,53^. Based on the observed effects of peri-LC_NPY_ activation on LC_NE_ neuronal excitability, we next hypothesized that local NPY signaling could represent an endogenous mechanism responsible for the regulation of anxiety in novel and/or anxiogenic environments. To test this hypothesis, we assessed the behavioral implications of peri-LC_NPY_ stimulation *in vivo*. *NPY-cre* mice were bilaterally injected with a Cre-dependent excitatory DREADD (AAV-hSyn-DIO-hM3D(Gq)-mCherry), or a vector expressing a control fluorescent protein (AAV-hSyn-DIO-mCherry) in the LC. After a period allowing for virus expression (5 weeks), mice were administered C21 (2mg/kg, i.p.) and were subjected to the EPM task (Fig. 4A). In hM3Dq mice, C21 increased the time spent in the open arm of the maze (mCherry, 8.1%, hM3Dq, 11.9% of total exploration time) and the frequency of open arm visits (mCherry, 10.0, hM3Dq, 14.4), indicating a reduction in baseline anxiety levels. In agreement, hM3Dq mice displayed decreased anxiety index (mCherry, 0.80, hM3Dq, 0.74; Fig. 4B). This anxiolytic effect occurred in the absence of C21 effects on general locomotion (Distance moved: mCherry 9.6m, hM3Dq, 10.3m; Fig. S7A). Likewise, controlling for possible off-target effects of C21, in a separate cohort of wild-type C57BL/6 mice, we confirmed that there were no gross behavioral changes (EPM performance, locomotion) after systemic C21 administration (Fig. S7C).

**Figure 4.**
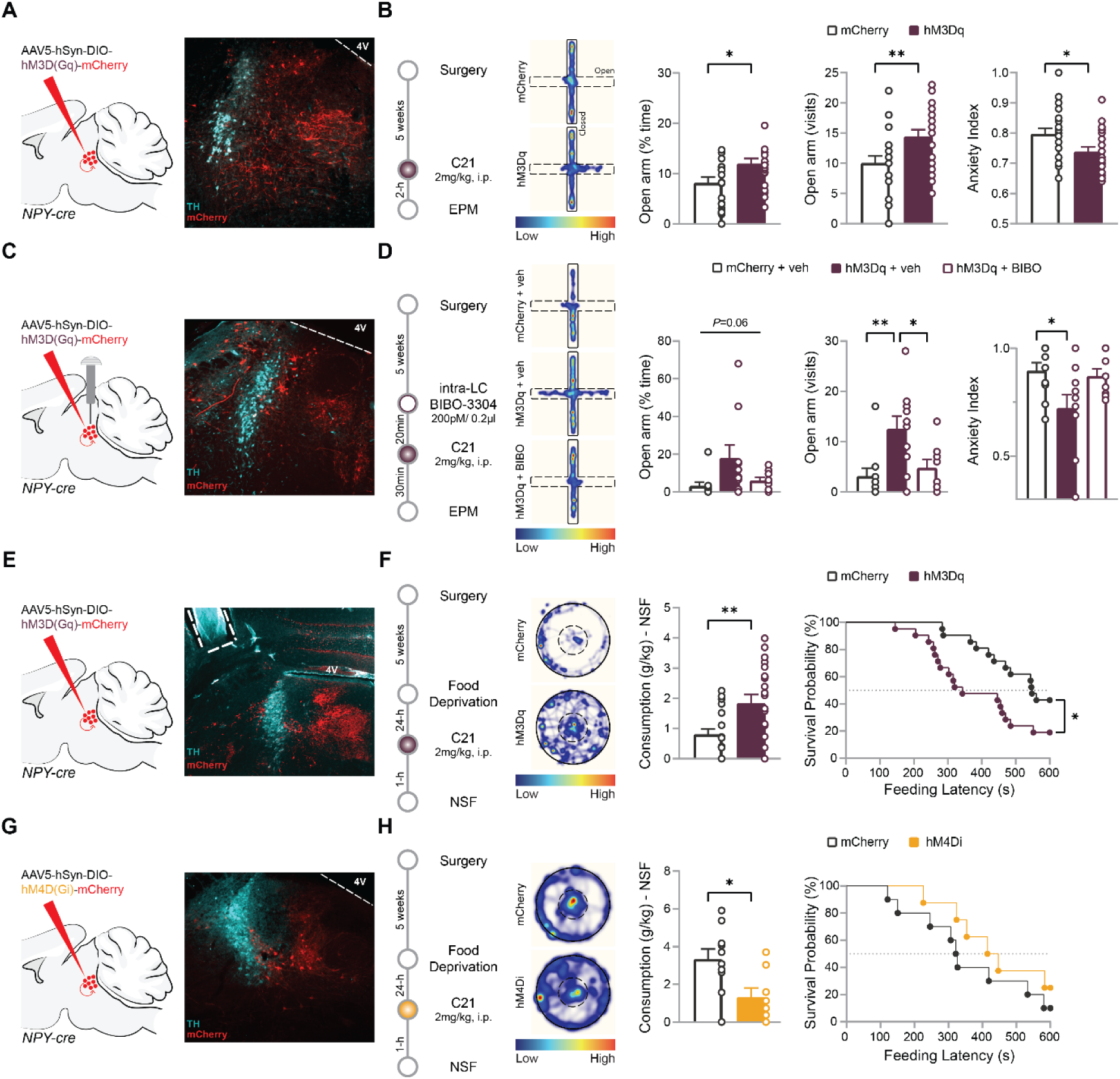
Peri-LC_NPY_ neurons bidirectionally regulate anxiety-like behavior. Experimental design for *in vivo* chemogenetic manipulations: schematic of sagittal mouse brain, depicting the location of bilateral virus injections used to drive hM3D(Gq) (A, C, E), hM4D(Gi) (G) or control vector expression in *NPY-cre* mice. Representative examples of virus spread, and cannula placement are included. B) After 5 weeks allowing for virus expression, mice were systemically administered the DREADD agonist compound 21 (C21, 2mg/kg, i.p.). Two hours after C21 injection, mice were subjected to the elevated plus maze (EPM) task. The hM3Dq group showed increased time spent in the open arms of the EPM (mCherry, 8.1%, hM3Dq, 11.9% of total exploration time; unpaired t-test, t(40)=2.42, *P*=0.020) and increased open arm entries (mCherry, 10.0, hM3Dq, 14.4; unpaired t-test, t(40)=2.71, *P*=0.010) *vs.* mCherry controls. Furthermore, in hM3Dq mice, C21 decreased anxiety index compared to mCherry controls (mCherry, 0.80, hM3Dq, 0.74; unpaired t-test, t(40)=2.31, *P*=0.026. D) After 5 weeks allowing for virus expression, cannula-assisted, bilateral, intra-LC micro-infusions of BIBO-3304 (200nM / 0.2µl / side) took place, 20min before systemic C21 administration (2mg/kg, i.p.). Thirty minutes after C21 injection, mice were subjected to the EPM task. A trend for main group effects on time spent in the open arms of the EPM was seen (mCherry & veh, 3.2%; hM3Dq & veh, 18.0%; hM3Dq & BIBO, 5.9% of total exploration time; 1-way ANOVA, F(2,25)=3.16, *P*=0.060). We detected main group effects for the number of visits to the open arms (mCherry & veh, 3.1; hM3Dq & veh, 12.5; hM3Dq & BIBO, 4.7; 1-way ANOVA, F(2,25)=6.16, *P*=0.007). This was due to hM3Dq & veh mice displaying significantly more entries *vs.* the other two groups (*P*=0.008 *vs.* mCherry & veh; *P*=0.042 *vs.* hM3Dq & BIBO). Likewise, we observed main group effects for anxiety index (mCherry & veh, 0.89; hM3Dq & veh, 0.72; hM3Dq & BIBO, 0.87; 1-way ANOVA, F(2,25)=3.80, *P*=0.036), driven by decreased anxiety index in hM3Dq & veh group *vs.* mCherry & veh controls (*P*=0.042). F, H) After 5 weeks allowing for virus expression, food deprived (24-h) mice were administered C21 (2mg/kg, i.p.). One hour after C21 injection, mice were subjected to the novelty suppressed feeding (NSF) task. F) hM3Dq mice showed increased food intake *vs.* mCherry controls (mCherry, 0.80 g/kg; hM3Dq, 1.84 g/kg; unpaired t-test, t(40)=3.03, *P*=0.004. Furthermore, C21 decreased feeding latency in the hM3Dq group (mCherry, 548s; hM3Dq, 343s; Mantel-Cox, χ^2^(1)=5.52, *P*=0.019). H) hM4Di mice consumed significantly less food *vs.* mCherry controls (mCherry, 3.33 g/kg; hM4Di, 1.32 g/kg; unpaired t-test, t(16)=2.66, *P*=0.017). No effects of C21 on feeding latency were seen (mCherry, 324s; hM4Di, 431s; Mantel-Cox, χ^2^(1)=1.24, *P*=0.265). B, D, F, H: Representative spatial location heatmaps show time spent exploring the arenas. B, F) Veh, N=21; hM3Dq N=21. D) mCherry & veh, N=10; hM3Dq & veh, N=10; hM3Dq & BIBO-3304, N=8. H) mCherry, N=10; hM4Di, N=8. Data depicted as mean ± SEM. * *P*<0.05. ** *P* <0.01.

Our findings suggest that NPY release by peri-LC_NPY_ neurons induces anxiolysis. We next assessed whether, in accordance with our electrophysiological data (*cf.,* Fig. 3H), this effect was mediated by local LC Y1R activation. To address this, we next performed an EPM experiment combining peri-LC_NPY_ chemogenetic activation with local, intra-LC Y1R antagonism *in vivo*. For this, *NPY-cre* mice expressing the excitatory DREADD (hM3D(Gq)) or mCherry control virus in peri-LC_NPY_ cells were bilaterally equipped with intra-cranial cannulas for LC-targeted administration of the selective Y1R antagonist BIBO-3304. After a period allowing for virus expression (≥5 weeks), we micro-infused mice with BIBO (200pmol/ 0.2µl/ side) or vehicle in the LC, before systemically administering C21 (2mg/kg, i.p) and then placed mice in the EPM (Fig 4C, D). As before, vehicle-infused hM3Dq mice (N=10), exhibited reduced anxiety-like behaviors *vs.* vehicle-infused mCherry controls. Notably, the anxiolytic effect of peri-LC_NPY_ chemogenetic stimulation was abolished in hM3Dq-expressing mice pre-treated with the Y1R antagonist. Particularly, hM3Dq+veh mice spent more time exploring the open arms of the EPM in comparison to the other two groups (mCherry+veh, 3.2%; hM3Dq+veh, 18.0%; hM3Dq+BIBO, 5.9% of total exploration time), although this fell short of statistical significance. Furthermore, a main group effect was observed for open arm entries (mCherry+veh, 3.1; hM3Dq+veh, 12.5; hM3Dq+BIBO, 4.7), driven by increased number of visits as seen in the hM3Dq+veh group compared to other groups (Fig 4D). Lastly, we detected a main group effect for anxiety index (mCherry+veh, 0.89; hM3Dq+veh, 0.72; hM3Dq+BIBO, 0.87), driven by decreased anxiety index in hM3Dq+veh group *vs.* mCherry+veh controls (Fig 4D). Unlike in the prior, non-cannulated experiment (*cf.*, Fig 4A, B), we here observed group effects on general locomotor activity (Distance moved: mCherry+veh, 5.3m; hM3Dq+veh, 10.4m; hM3Dq+BIBO, 6.8m; Fig. S7B). Nonetheless, our findings confirmed a causal link between peri-LC_NPY_ activity and the regulation of anxiety-like behaviors, where local Y1R-mediated NPY signal tempers LC_NE_ firing and promotes anxiolysis.

Our data suggest that peri-LC_NPY_-mediated input to the LC sufficiently drives anxiolysis in the EPM. To further corroborate this we evaluated the role of peri-LC_NPY_ neurons in another behavioral assay, namely the novelty-suppressed feeding (NSF) test, which assesses hyponeophagia, the suppression of food intake due to exposure to a novel environment, as a proxy for anxiety^54^. For this, we used the aforementioned cohort of *NPY-cre* mice with Cre-dependent excitatory DREADD (hM3Dq), or control virus (mCherry) in peri-LC_NPY_ neurons. Mice were food deprived for 24-h, followed by systemic C21 administration (2mg/kg, i.p.) and assessed in the NSF task (Fig. 4E). In hM3Dq mice, C21 decreased the latency to initiate consumption of a familiar food source placed at the center of an open field arena (mCherry, 548s; hM3Dq, 343s), indicating reduction in anxiety levels. In support, hM3Dq mice spent more time at the center of the arena (Duration Center: mCherry, 7.7%; hM3Dq, 13.4% of total exploration time), where they consumed significantly more food *vs.* mCherry controls (NSF intake: mCherry, 0.80 g/kg; hM3Dq, 1.84 g/kg; Fig. 4F, Fig. S7D). Controlling for effects of peri-LC_NPY_ chemogenetic stimulation on general consummatory behavior, we observed no group differences on food consumption in a familiar environment (Home-cage intake: mCherry, 6.76 g/kg; hM3Dq, 7.35 g/kg; Fig. S7D). No effects of C21 on general locomotor activity at the NSF were seen (Distance moved: mCherry 41.5m, hM3Dq, 42.7m; Fig. S7D), validating the specificity of the anxiolytic effects of peri-LC_NPY_ activation.

Overall, our data support that peri-LC_NPY_ activation sufficiently drives anxiety relief, however, whether it is also required for anxiolysis remains unresolved. To address this, we examined the effects of chemogenetic peri-LC_NPY_ inactivation on EPM and NSF tasks. For this, we made use of a Cre-dependent AAV (AAV-hSyn-DIO-hM4D(Gi)-mCherry) that drives the expression of an inhibitory DREADD. First, using electrophysiological recordings, we confirmed that hM4Di activation leads to peri-LC_NPY_ cells silencing (Fig. S8A, B). In particular, bath application of C21 (2µM), resulted in reduced peri-LC_NPY_ spontaneous firing, as recorded at current clamp mode. Then, in an independent cohort of *NPY-cre* mice, we bilaterally injected the inhibitory DREADD (AAV-hSyn-DIO-hM4D(Gi)-mCherry), or a control virus (AAV-hSyn-DIO-mCherry) in the LC. After a period allowing for virus expression (5 weeks), mice were food deprived (24-h), were administered C21 (2mg/kg, i.p.) and were then subjected to the NSF task, as described above (Fig. 4G). In hM4Di mice, C21 administration reduced food intake at the NSF arena, as compared to mCherry controls (NSF intake: mCherry, 3.33 g/kg; hM4Di, 1.32 g/kg), indicative of peri-LC_NPY_ inhibition-triggered anxiogenesis (Fig. 4H). This effect on food intake was specific for anxiogenic contexts, as it did not occur when food was available at the home-cage (Home-cage intake: mCherry, 8.62 g/kg; hM4Di, 8.97 g/kg, Fig. S8C). Peri-LC_NPY_ inhibition by C21 did not affect the latency to initiate food consumption in the NSF arena (mCherry, 324s; hM4Di, 431s), nor the time spent at its center (Duration Center: mCherry, 13.1 %; hM4Di, 13.0% of total exploration time, Fig. 4H, Fig. S8C). As we observed above for NSF, C21 did not alter general locomotion in hM4Di mice (mCherry, 10.7m; hM4Di, 10.8m) (Fig. S8C). Notably, peri-LC_NPY_ inhibition did not affect performance on the EPM (Fig. S8D), indicating differential involvement of peri-LC_NPY_ inhibition in these two distinct anxiety tests. Together, our data support bidirectional control of (certain) anxiety-like behaviors by peri-LC_NPY_ neurons, highlighting a novel endogenous mechanism for the promotion of adaptive responses to challenging environments.

## Discussion

Despite decades of research on the basic mechanisms underlying adaptive stress reactivity, our knowledge of the relevant neural circuitries, and molecular substrates remains incomplete. The LC_NE_ system is the brain’s “first responder” to stress and primary coordinator of the global neural processes underlying flight-or-fight^7,9,55^. Therefore, we here reasoned that elucidating the endogenous systems that curb LC function, could provide us with a mechanistic understanding on how to promote adaptive responses to challenging environments. For this, we examined the endogenous mechanisms underlying NPY-mediated LC neuromodulation. Collectively, we here demonstrate that a previously uncharacterized population of NPY-expressing cells, situated in the pericoerulean region, provides direct, Y1R-mediated NPY tone onto LC noradrenergic neurons, tempering their activity. Under approach-avoidance conflicts, peri-LC_NPY_-mediated neuromodulation of the LC efficiently lowers anxiety levels, via Y1R-dependent mechanisms. Conversely, suppressing local peri-LC_NPY_ neuronal activity results in anxiogenesis. Together, our findings highlight a role for peri-LC_NPY_ neurons as key mediators of LC function and delineate a novel endogenous circuit for the regulation of (trait) anxiety.

We report that the large majority of peri-LC_NPY_ neurons are situated medially to the LC nuclear core, within a region that is rich both in noradrenergic dendritic fibers and diverse axon terminals targeting them. Convergent signals within this area, mediated by fast neurotransmission or slow peptidergic input^12,51,56–58^, are proposed to coordinate LC engagement in arousal and under stress. Thus, peri-LC_NPY_ neurons are ideally situated to fine-tune LC_NE_ activity levels when required. Our neuroanatomical tracing data support this notion, as we provide evidence that peri-LC_NPY_ neurons project to the LC dendritic zone, and form functional contacts within the region, and with LC_NE_ neurons themselves. Notably, using viral anterograde tracing we did not detect peri-LC_NPY_ neuronal efferents in other primary LC terminal fields. However, given that peri-LC_NPY_ cells comprise a relatively small population, we cannot exclude that unbundled projection fibers remained undetected in our preparations. Furthermore, our retrograde tracing data revealed several novel brain regions that project to the LC, indicating that peri-LC_NPY_ cells are not the sole source of NPY in the region. Future studies are required to unravel non-LC NPY contributions to the neuromodulation of the LC noradrenergic system.

In contrast to earlier studies in rats^21,28,59^, we here report that peri-LC_NPY_ neurons constitute a distinct, predominantly non-noradrenergic neuronal population. Although species differences might contribute to the observed discrepancy, it is also plausible that methodological limitations previously overestimated NPY-NE colocalization in the LC proper. Using more advanced techniques, recent single-cell and single-nucleus RNA sequencing data corroborated lack of NPY-expressing noradrenergic neurons, both in the mouse^31,60^ and in the human^61^ LC. Notably, similar to our observations, NPY transcripts were detected outside the LC nuclear core^31^, further validating the existence of a previously undetected discrete pool of NPY-expressing neurons populating the mouse pericoerulean space.

We here show that NPY Y1R activation directly affects LC_NE_ activity. Y1Rs are preferentially localized at post-synaptic sites^30,62^, ideally placing them to regulate LC_NE_ intrinsic excitability. Notably, the effects we detect here resemble that of exogenous NPY application in the lateral amygdala^46^. In particular, in amygdalar projection neurons, pharmacologically applied NPY had a hyperpolarizing effect via Y1 (and not Y2) receptors, by activation of inward rectifying potassium channels^46^. Following peri-LC_NPY_ chemogenetic activation, we did not detect changes in LC_NE_ membrane resistance, calculated near resting membrane potential. However, we cannot exclude differences in potassium channel-regulated membrane resistance at more positive potentials, which we did not assess here, as in our experiment those coincided with action potential-driven active conductances. Likewise, we did not specifically address the contribution of GIRK-mediated currents on C21-induced hyperpolarization of LC_NE_ cells, thus we cannot exclude their involvement in the observed effects.

We here demonstrate that the anxiety-relieving effects of local NPY release in the LC are mediated via Y1Rs, linking our electrophysiological observations with the behavioral outcomes of peri-LC_NPY_ chemogenetic activation. Interestingly, our data are in conflict with an earlier study in rats^25^, where pharmacologically applied NPY within the LC vicinity resulted in reduced anxiety via Y2Rs. In particular, Kask and colleagues (1998) showed that infusion of NPY or an agonist with affinity for Y2/Y5 receptors in the pericoerulean space had an anxiolytic effect in rats performing the EPM task. On the contrary, a singular low dose of a Y1R/Y5R agonist did not produce this effect. These findings seem at odds with our observations that implicate Y1Rs in the regulation of anxiety-like behaviors following chemogenetically-evoked release by local NPY sources. There are obvious key differences between the two studies. First, exogenously applied NPY can induce widespread recruitment of Y2R-containing efferents outside the LC dendritic zone, with extra-LC contributions accounting for the observed behavioral effects. Instead, we here assess recruitment of local endogenous NPY sources, possibly relying on proximity between pre-and post-synaptic targets, as seen for NE^63^, resulting in targeted effects on LC_NE_ intrinsic excitability and the ensuing Y1R-dependent anxiolysis.

Moreover, pharmacological dose chosen could contribute to the apparent disparity between the two studies. Indeed, we here report that the pharmacological effects of exogenously applied NPY on LC_NE_ activity are dose-dependent. In fact, bath-applied NPY modulates LC_NE_ neurons in opposite directions (hypo-*vs.* hyper-excitability), via distinct NPY receptors. Particularly, low NPY dose mimicked the effects of DREADD-induced activation of peri-LC_NPY_ neurons, with both depending on postsynaptic Y1Rs. High NPY dose led to opposite effects, via a network mechanism that relied on Y2Rs and synaptic input onto LC_NE_ cells. While we did not specifically test this, it is possible that presynaptic Y2Rs, expressed on peri-LC_NPY_ cells, play an auto-regulatory role^64^, in which they moderate peri-LC_NPY_ signaling, leading to disinhibition of LC_NE_ neurons and increased LC_NE_ firing frequency. Exposure to severe stress might require such a mechanism when rapid stress response is the preferable and adaptive choice, and Y2R-mediated signaling is shown to fulfil this role^65–68^. On the other hand, in mildly anxiogenic environments, prioritizing the regulation of the initial stress response by Y1Rs might preclude the development of maladaptive behavioral patterns. Indeed, in several brain regions other than the LC, Y1R activation results in anxiolysis^17,19,69–71^. Collectively, these results highlight the intricate outcomes of NPY-mediated neuromodulation of the LC_NE_ system, where NPY tone can fine-tune LC_NE_ activity towards both hypo-and hyper-excitation. Which physiological conditions prompt each requirement *in vivo* remains to be determined.

In the present study, we addressed the role of peri-LC_NPY_ neurons in the regulation of basal anxiety levels. However, their involvement in modulating behavior after exposure to stress remains unclear. Stress elevates NPY mRNA expression in the LC^72^, which parallels our findings on stress-induced engagement of peri-LC_NPY_ neurons, as seen in increased firing frequency and cFos-indicated neuronal activation. Moreover, stress enhances LC Y1R and Y2R mRNA levels^73^, further indicating recruitment of the local NPY system during adversity. In rats, signal intranasal administration of NPY limits the development of hyperarousal and anxiety-like behaviors seen after exposure to stress^72^. Notably, this is accompanied by attenuated stress-induced increases in LC TH expression, which could reflect curbing of LC function. It is possible that, in a similar manner, peri-LC_NPY_ recruitment and local NPY release contribute to LC silencing and the promotion of adaptive responses under stress.

Combining *in vivo* chemogenetic manipulations with behavioral assessment, we here showed that modulation of peri-LC_NPY_ neuronal activity exerts bidirectional control over anxiety-like manifestations. Particularly, peri-LC_NPY_ stimulation increased exploration/approach in the EPM and NSF tasks, in agreement with peri-LC_NPY_-mediated silencing of LC_NE_ cells, and presumably, reduced NE release in terminal fields, promoting anxiolysis. Likewise, we demonstrate that peri-LC_NPY_ inhibition, which is expected to relieve an NPY brake on LC_NE_ neurons, led to anxiogenesis at the NSF. Notably, we report the lack of effect of *in vivo* peri-LC_NPY_ inhibition in the EPM, implying divergent requirements for peri-LC_NPY_ function in each task conditions. Of note, in the NSF, but not the EPM, mice were subjected to prior food deprivation, which is shown to increase LC neuronal activity during food approach^53^. Increased LC engagement could set the stage for enhanced peri-LC_NPY_ recruitment upon fasting. Thus, our NSF data might reflect an adaptive mechanism aimed to counteract increased LC_NE_ tonic activity-induced avoidance/ aversion^12,53^, in favor of exploration of / foraging in new -and potentially unsafe environments-under conditions of limited food availability. In accordance with this notion, whereas peri-LC_NPY_ activity levels influenced food-drive and consumption in the anxiogenic NSF task, it left consummatory behavior in the familiar (safe) context of the home-cage unaltered.

In the current study, we performed cell type-specific dissection of a novel circuitry mediating anxiety relief. In particular, we provide mechanistic data for the anxiolytic effects of a previously unidentified NPY-expressing neuronal population. Using combinatory approaches, we showcase anatomical, electrophysiological and behavioral evidence that peri-LC_NPY_ neurons exert neuromodulatory control over the LC_NE_ system, promoting its adaptive engagement in arousing environments, and the maintenance of low baseline anxiety levels. Collectively, these data expand our understanding of how endogenous peptidergic influences regulate the brain’s stress systems and highlight peri-LC_NPY_ neurons as a possible new target for the modulation of anxiety-like behaviors.

## Methods

### Animals

In all experiments naïve adult male or female mice were used (20–35 g, ≥4 weeks). C57BL/6J (Charles River, France), *NPY-Cre* (Jax #027851) and *Ai14* (Jax #007914) mice were bred in house after purchasing founders from the original breeding colonies. *NPY-cre:Ai14* were bred in house. All mice were group-housed (2–5 per cage) unless otherwise specified. Mice were housed in a 12:12 light/dark cycle (lights on at 07:00 a.m.) at 22 ± 2 °C (60–65% humidity). Unless otherwise specified, animals had access to ad libitum water and lab chow. Experiments were approved by the Animal Ethics Committee of Utrecht University and the Dutch Central Authority for Scientific Procedures on Animals (CCD) and were conducted in agreement with the Dutch law (Wet op de Dierproeven, 2014) and European regulations (Guideline 86/609/EEC).

### Stereotactic surgeries

Mice (≥ 5 weeks at the time of surgery) were anesthetized with ketamine (75 mg/kg i.p.; Narketan, Vetoquinol) and dexmedetomidine (1 mg/kg i.p.; dexdomitor, Vetoquinol). Lidocaine (0.1 ml; 10% in saline; B. Braun) was injected under the skull skin and eye ointment cream (CAF, Ceva Sante Animale B.V.) was applied. Animals were then fixed on a stereotactic frame (UNO B.V. model 68U801 or 68U025), where they kept on a heat pad (33 °C) during surgery. For cannula implantations, the skull surface was scratched with a scalpel and phosphoric acid (167-CE, ultra-Etch, ultradent, USA) was applied for 5 min to roughen the surface at the start of surgery. Viral infusions were done using a 31 G metal needle (Coopers Needleworks) attached to a 10 µl Hamilton syringe (model 801RN) via flexible tubing (PE10, 0.28 mm ID, 0.61 mm OD, Portex). The Hamilton syringe was controlled by an automated pump (UNO B.V., model 220). Injections were done bilaterally at 250 nl per side, at an injection rate of 100 nl/min. The injector was gradually retracted during the last minute of a 10 min long diffusion period. Next, the skin was sutured (V926H, 6/0, VICRYL, Ethicon) and animals were administered the dexmedetomidine antagonist atipamezole (50 mg/kg; Atipam, Dechra, s.c.), carprofen (5 mg/kg, Carporal, s.c.) and 1 ml of saline (s.c.). Mice were left to recover on a heat plate (36 °C) while being monitored and moved to the housing stables when fully awake. Carprofen (0.025 mg/L) was provided in the drinking water during the first post-operative week. Post-surgery, animals were single-housed before rejoining their cage-mates at three days post-op. Animals with cannulas were single-housed for the remainder of the experiment.

For *ex vivo* optogenetic or chemogenetic-assisted electrophysiology experiments and ELISA, *NPY-cre* mice were bilaterally injected with rAAV5-Syn-FLEX-CoChR-GFP (4.4*10^12 gc/ml; UNC Vector Core) or rAAV5-hSyn-DIO-hM3D(Gq)-mCherry (4.2*10^12 gc/ml; Addgene) in the Locus Coeruleus (LC, AP: -5.45 mm; ML: ± 1.59 mm; and DV: -3.96 mm from bregma) under a 10 degree angle. All animals were allowed to recover for a minimum of four weeks before brain collection for electrophysiological recordings and other biochemical assays.

For *in vivo* chemogenetic experiments, *NPY-cre* mice were bilaterally injected with rAAV5-hSyn-DIO-hM3D(Gq)-mCherry, AAV5-hSyn-DIO-hM4D(Gi)-mCherry or AAV5-hSyn-DIO-mCherry (all viruses: 4.2*10^12 gc/ml; Addgene) in the LC (AP: -5.45 mm; ML: ± 1.59 mm; DV: -3.96 mm or AP: -5.3 mm; ML: ± 1.95 mm; DV: 3.5 mm from bregma) under a 10 degree angle. All animals were allowed to recover for five weeks before participating in behavioral assays.

For cannula implantations, *NPY-cre* mice were bilaterally implanted with a guide cannula (4 mm; C315GMN/Spc; Bilaney) above the LC (AP: -5.35 mm; ML: ± 1.63 mm; and DV: -3.76 mm or AP: - 5.35 mm; ML: ± 1.77 mm; and DV: -3.86 mm from bregma) under a 10 degree angle. The cannulas were secured by adding a layer of adhesive luting cement (C&B Metabond; Parkell, Edgewood, NY, USA) around them. Viral infusions were performed immediately after cannula placement, by inserting the injector through the guide, and lowering it to DV -3.96 mm. After infusions, dummy internal injectors (4 mm; C315FD/Spc) were locked onto the guides to avoid blockage, and remained there until experiment completion. All animals were allowed to recover for five weeks before participating in behavioral assays.

### Behavioral assays

All behavioral manipulations and tests were performed during the light phase, between 12-6pm. Only male mice were included in the behavioral assays described below. Animals were transported to the behavioral room at least 2 weeks before the start of the experiment, for acclimatization. Real-time data collection and offline analysis for EPM and NSF tasks were done with Ethovision video tracking (version XT 9; Noldus).

#### Foot-shock stress

Exposure to foot-shock stress was conducted in operant chambers (29.53 × 24.84 × 18.67 cm, Med Associates) modified for foot shock delivery, via the grid floor. The chambers were contained in a soundproofing box. Mice were placed in the chamber and given 5 min of habituation to the novel environment. A total of 20 foot-shocks (1 s duration, 0.3 mA intensity) were delivered with an interval of 60s for the next 20 min of the session. The last 5 min of the session (30min of total duration) were shock-free. Control (no-stress) mice were exposed to the operant chamber (novel environment) in absence of electrical foot-shocks. After the session, all mice were transferred back to their home-cage.

#### Open Field

The open field (OF) task was performed using a standard arena (round, diameter 80 cm). Light intensity in the center of the arena was 25 Lux. At the start of the session, mice were placed next to the walls of the arena and allowed to freely explore for 10 min. Behavior was scored for the following variables: time spent in the center or wall zone (% of total exploration time); frequency of visits to the center or wall zone; and total distance moved. The OF arena was cleaned with 70% ethanol solution in-between animals.

#### Elevated Plus Maze

The elevated plus maze (EPM) task was performed using a standard apparatus (arm length: 65 cm × 65 cm) elevated at 65 cm above ground, equipped with 15 cm high walls delimiting the enclosed arms. Light intensity in the center of the maze was 40 Lux. At the start of the session, mice were placed at one of the closed arms, facing the EPM center, and allowed to freely explore the maze for 10 min. Behavior was scored for the following variables in the first 5 minutes of the EPM: time spent in the open or closed arms (% of total exploration time); frequency of visits to the open or closed arms; total distance moved; and anxiety index^36^, calculated as: 1 - [(time in OA / (time in OA + time in CA)) + (entries to OA / (entries to OA + entries to CA)) / 2], where values closer to 1 delineate increased anxiety. EPM was cleaned with 70% ethanol solution in-between animals.

#### Novelty Suppressed Feeding

The novelty suppressed feeding (NSF) task was performed using the open field arena as described above. A metal disk containing standard lab chow (1 pellet) was firmly affixed at the center of the arena. The pellet itself was attached to the disk, so that the animals could not move it during the test. At the start of the session, mice were placed next to the walls of the arena and allowed to freely explore for 10 min. Behavior was scored for the following variables: time spent in the center or wall zone (% of total exploration time); frequency of visits to the center or wall zone; and total distance moved. In addition, latency to initiate feeding and food consumption in the arena, calculated as: food pellet weight at start - end of session / mouse weight (g/kg), were recorded. A different disk, and chow pellet were used per mouse, and the NSF arena was cleaned with 70% ethanol solution in-between animals.

### *In vivo* chemogenetic experiments

All animals were habituated to restrain for intraperitoneal (i.p.) injections with at least 3x handling sessions that included “sham” injections before the start of the experiments. For intracranial infusions, mice were extensively handled for at least a week (1x day) for familiarization with infusion procedures (dummy removal, injector/ tubing plug-in), before the start of the experiments. For all experiments, only mice with confirmed viral expression within the LC and, when applicable, correct cannula placement after histological control were included in data analysis.

For peri-LC_NPY_ stimulation experiments, two separate batches of *NPY-cre* mice were used, with 2 months interval between experiments. Both cohorts (Batch 1: mCherry, N=11; hM3D(Gq), N=11; Batch 2: mCherry, N=10; hM3D(Gq), N=10) displayed similar performance, thus data were pooled.

For intra-LC Y1 antagonism, two separate batches of *NPY-cre* mice were used, with 2.5 months interval between experiments. Both cohorts (Batch 1: mCherry + veh, N=4; hM3D(Gq) + veh, N=5; hM3D(Gq) + BIBO, N=5; Batch 2: mCherry + veh, N=6; hM3D(Gq) + veh, N=5; hM3D(Gq) + BIBO, N=3) displayed similar performance, thus data were pooled.

#### Elevated Plus Maze

For peri-LC_NPY_ stimulation or inhibition experiments, all mice were administered C21 (2mg/kg, i.p.). One hour after injection, animals were exposed to a novel environment (MED apparatus, as described above) for 30min, before being transferred to a clean cage, where they remain single-housed for the rest of the experimental procedures. Thirty minutes later, mice were subjected to the EPM test, before returning to their home-cage.

For intra-LC Y1 antagonism experiments, internal guides (1 mm projection; C315IMN/Spc; Bilaney) attached to tubing (PE10, 0.28 mm ID, 0.61 mm OD, Portex) were bilaterally inserted in the cannulas. The tubing was attached to 10 µl Hamilton syringe (model 801RN), controlled by an automated pump (UNO B.V., model 220). While the animals were in their home cage, 200pmol BIBO-3304 or vehicle was infused in a volume of 200 nl (1% DMSO, 99% PBS) in each hemisphere, at a rate of 100 nl/min. The injector was left in place for an additional minute. Twenty minutes after intra-cannula infusions, animals were administered C21 (2mg/kg, i.p.). Thirty-five minutes after C21 injections, mice were subjected to the EPM task, as described above.

#### Novelty suppressed feeding

For peri-LC_NPY_ stimulation or inhibition experiments, mice were subjected to the NSF task, 48 hours after participating in the EPM and following 24-h of food deprivation. All mice were administered C21 (2mg/kg, i.p.) and NSF took place one hour after injection. Immediately after the end of the session, mice were returned to their home-cage where food (chow pellet) intake was recorded for 5-min.

### Patch-clamp electrophysiology

Animals were anesthetized with pentobarbital (Euthasol 20%, 0.1ml, i.p.) between 8.30 a.m. and 10 a.m. and transcardially perfused with ice-cold carbogenated (95% O2, 5% CO2) slicing solution containing (in mM): choline chloride 92; ascorbic acid 10; CaCl2 0.5; glucose 25; HEPES 20; KCl 2.5; N-Acetyl L Cysteine 3.1; NaHCO3 25; NaH2PO4 1.2; NMDG 29; MgCl2 7; sodium pyruvate 3; Thiourea 2. Brains were quickly extracted, placed on a vibratome (1200 VTs, Leica) and sliced in the coronal plane at 250 μm thickness, in ice-cold slicing solution. Slices recovery was performed for 30 min at 36 °C in carbogenated solution of identical composition. Thereafter, slices were maintained at room temperature in carbogenated incubation solution containing (in mM): ascorbic acid 3; CaCl2 2; glucose 25; HEPES 20; KCl 2.5; NaCl 92; NaHCO3 20; NaH2PO4 1.2; NMDG 29; MgCl2 2; sodium pyruvate 3 and Thiourea 2. During recordings, slices were immersed in artificial cerebrospinal fluid (ACSF) containing (in mM): CaCl2 2.5; glucose 11; HEPES 5; KCl 2.5; NaCl 124; NaHCO3 26; NaH2PO4 1; MgCl2 1.3 and were continuously superfused at a flow rate of 2 ml min^-1^ at 28-30 °C.

Peri-LC_NPY_ or LC_NE_ neurons were patch-clamped using borosilicate glass pipettes (2.7–4.5 MΩ; glass capillaries, GC150-10, Harvard apparatus, UK), under a TH4-200 Olympus microscope (Olympus, France). For voltage or current clamp recordings, signals were amplified and digitized using a HEKA EPC-10 patch-clamp amplifier (HEKA Elektronik GmbH). Data were acquired using PatchMaster v2×90.2 software. Access resistance was continuously monitored with a − 4 mV step delivered at 0.1 Hz. Experiments were discarded if the access resistance increased by more than 20% during the recording. All electrophysiological measures are recorded with a 10 s inter sweep interval (0.1 Hz).

#### Intrinsic excitability

Recordings were made in a potassium gluconate-based internal containing (in mM): Kglu 139; HEPES 10; EGTA 0.2; creatine phosphate 10; KCl 5; Na2ATP 4; Na3GTP 0.3; MgCl2 2. Upon break-in, cells were kept at -50mV for 10 min prior to the onset of current-clamp recordings. To assess passive membrane properties and cell firing patterns neurons were subjected to 17 consecutive current steps of 800 ms length, starting from −150 to +250 pA, with a 25 pA inter-step increment.

For stress effects on peri-LC_NPY_ or LC_NE_ excitability, LC-containing slices from *NPY-cre:Ai14* mice were obtained 30 min after foot-shock stress or exposure to the novel environment (no-stress controls). LC_NPY_ cells were identified by tdTomato expression under the microscope. LC_NE_ neurons were identified based on location and morphological characteristics.

For *ex vivo* chemogenetic experiments, effects of bath-applied C21 were examined in LC-containing slices from *NPY-cre* mice at ≥5 weeks after DIO-hM3D(Gq) injection. Recordings were made in continuous perfusion of C21 (2 µM) or vehicle (0.1% DMSO). When applicable, slices were pretreated with synaptic blockers (in µM: CNQX 10, D-AP5 50, picrotoxin 100, CGP-54626 10), Y1R (BIBO-3304, 1 µM) or Y2R (BIIE-0246, 1 µM) antagonists for 10 min before being transferred in ACSF containing C21 or vehicle and the correspondent antagonist mix. All slices for chemogenetic experiments were controlled for virus expression at the end of recordings. Only data from slices with strong hM3D(Gq) expression within the peri-LC were taken along for analysis.

For NPY effects on LC_NE_ excitability, LC-containing slices were obtained from wild-type C57BL/6 mice. Recordings were made in continuous perfusion of NPY (30 or 300 nM) or vehicle (0.1% DMSO). When applicable, slices were pretreated (10 min) with synaptic blockers as described above, before being transferred in ACSF containing NPY or vehicle.

For validation of DREADD constructs, effects of bath-applied C21 were examined in LC-containing slices from *NPY-cre* mice at ≥5 weeks after DIO-hM3D(Gq) or DIO-hM4D(Gi) injection. Peri-LC_NPY_ cells were identified by mCherry expression under the microscope. Peri-LC_NPY_ spontaneous activity was recorded before and after C21 (2 µM) bath application.

#### Peri-LC_NPY_ to LC_NE_ connectivity experiments

At ≥5 weeks after virus injection of FLEX-CoChR in the LC of *NPY-cre* mice, recordings were made in voltage clamp in a potassium gluconate-based internal solution containing (in mM): Kglu 139; HEPES 10; EGTA 0.2; creatine phosphate 10; KCl 5; Na2ATP 4; Na3GTP 0.3; MgCl2 2, at -50 mV for the detection of inward glutamatergic and outward GABAergic currents. Alternatively, recordings were performed with a cesium chloride-based internal solution containing (in mM): CsCl 139; HEPES 10; EGTA 0.2; creatine phosphate 10; NaCl 5; Na2ATP 4; Na3GTP 0.3; MgCl2 2; spermine 0.1, at -60 mV or +40 mV for detecting inward GABA-or outward NMDA-mediated currents, in correspondence. Responses to single optical pulse (470 nm, 1 ms, 1–2 mW) or trains of 5, 20 and 50 Hz delivered through the light path of the microscope powered by a light-emitting diode driver (LEDD1B; Thorlabs, Newton, NJ) were recorded. Connectivity was determined based on whether a neuron showed an opto-evoked synaptic response ≥ 5 pA, over an average of 10-20 sweeps. In a portion of recordings, internal solutions contained 3% biocytin (B4261, Sigma-Aldrich), for cell-filling and post-hoc identification. All slices for optogenetic experiments were controlled for virus expression at the end of recordings. Only data from slices with strong CoChR expression within the peri-LC were included in the connectivity analysis.

### Histology & Immunolabelling

All mice were anesthetized with pentobarbital (Euthasol, 0.1 ml, i.p.) and transcardially perfused with PBS followed by freshly-made ice-cold 4%PFA. Brains were extracted and post-fixated in 4% PFA overnight, before being transferred to an anti-freeze solution (30% sucrose) until they sank. Thereafter sections measuring 35 µm (colocalization studies) or 50 µm (histological control of virus targeting) were collected using a cryostat (Leica CM 1950) at -12°C and stored in PBS & 0.01% NaN_3_ until immunolabelling. Brains from animals participating in cannula experiments were post-fixated for 48-h and transferred to 10% sucrose before being embedded to a 10% sucrose/ 10% gelatin solution. Next, embedded brains were re-fixated in 10% sucrose/ 4% PFA solution overnight and stored in 30% sucrose until slicing. Sections (50 µm) were collected with a vibrotome (Leica VT 1000S) at room temperature (RT). In most cases, every 5^th^ consecutive section was collected for further processing, to ensure LC representation in the entire rostrocaudal axis. For immunohistochemical stainings, sections were washed in 1× PBS, and blocked (1-h, RT) in a solution containing 5% normal goat or donkey serum, 2.5% bovine serum albumin and 0.2% Triton X-100. Primary antibodies against NPY (Novus Biologicals, NBP1-46535), TH (LNC1, Millipore, MAB318), GABA (Sigma-Aldrich, A2052), and cFos (9F6, Cell Signaling Technology, 2250S) were incubated overnight at 4°C. Secondary antibodies (Alexa Fluor 488, 568 or 647, Invitrogen) were incubated for 2-h (RT). Slices were mounted in 0.2% gelatin and cover-slipped with DABCO antifading medium (Merck, 10981).

For NPY mapping studies, sections were imaged at 10x magnification using a confocal microscope (LSM 880, Zeiss). Tiling was performed to include the mediolateral space to LC, up to 1 mm from the LC proper. Images were processed with FIJI^74^ and an in-house macro was used for detection of tdT+ and TH-expressing cells. Location of NPY-expressing neurons was determined based on X (mediolateral) and Y (dorsoventral) coordinates as calculated against an ROI representing LC center of mass per image. Frequency distributions for NPY location in ML and DV axes were calculated for three different rostrocaudal ranges, namely-5.80mm to -5.60mm, -5.60mm to -5.40mm and -5.40mm to -5.25mm, and averaged over the corresponding images.

For co-localization studies, sections were imaged by confocal microscope at 20x or 40x magnification and z-stacks (≥ 8 images) were processed with FIJI. An in-house macro was used for detection of tdT^+^, TH^+^, and cFos^+^ cells, which overlaid detected ROIs in one channel over the second. For quantification of tdT^+^/cFos^+^ double-expressing cells the following inclusion criteria were used: 1) cFos channel intensity ≥50% of a positive, cFos-expressing “example” cell and 2) cFos channel intensity ≥ 150% of background.

### RNA-scope in situ hybridization assay and analysis

In situ hybridization (ISH) was performed as per manufacturer’s instructions (ACDBio, RNAscope Multiplex Fluorescent Reagent kit v2, 317621). Briefly, animals were transcardially perfused with sterile 4% PFA and post-fixated ovenight at 4 °C. Brains were transferred to 10% sucrose until they sank. This step was repeated with 20 and 30% sucrose. Brains were then embedded in optimal cutting temperature media (Tissue-Tek, VWR, The Netherlands) and placed in the cryostat at −20 °C for 1 h to equilibrate. Sections measuring 10 μm containing the LC were then mounted on SuperFrost Plus slides (VWR, The Netherlands) and allowed to dry at −20 °C for 2 h. For ISH, sections were pre-treated with hydrogen peroxide for 10 min before target retrieval at 99 °C for 5 min and treatment with Protease III for 30 min at 40 °C. Hybridization to probes against NPY (313321), Y1R (427021), Y2R (315951) or TH (317621) was carried out at 40 °C for 2 h. HRP signals against each channel (C1–C3) were then sequentially amplified and developed using TSA Vivid fluorophores (520, 570 and 650) at a dilution of 1:1500 or 1:3000. Positive-(3-plex PN 320881), and negative-control probes (3-plex PN 320871) were included in each experiment to assess sample RNA quality and optimal permeabilization conditions. In some cases, immunolabeling was combined with ISH. For this, sections were blocked with 0.1% BSA in TBS (30min, RT) before incubation of primary antibody against TH (1.5-h, RT). Sequential secondary HRP (30 min, RT) and TSA (10 min, RT) incubations were performed next. Sections were counterstained with DAPI for 30 s, coverslipped with ProLong Gold Antifade Mountant and allowed to dry overnight at room temperature. Confocal mages (40x magnification) were processed with FIJI and an in-house macro for detection of TH-or NPY-expressing cells.

### Data analysis, statistics and reproducibility

Data were analyzed with Noldus (Ethovision V9.0), GraphPad Prism (9.5.1), Igor Pro-8 (Wavemetrics, USA) and FIJI. Sample size was predetermined on the basis of published studies, experimental pilots and in-house expertise. Animals were randomly assigned to experimental groups. Compiled data are always reported and represented as mean +SEM, with single data points plotted (single cell for electrophysiology and single animal for behavioral experiments). Animals or data points were not excluded from analyses unless noted. When applicable, statistical comparisons were conducted using unpaired t-tests, and 1-way or 2-way repeated measure ANOVAs. When applicable post hoc comparisons were performed in case of significant main (interaction) effects. Normality distribution was confirmed with the Kolmogorov-Smirnov test and in case of violation non-parametric Mann-Whitney test was performed. Two-tailed testing was performed with significance level α set at 0.05.

## Acknowledgements

We thank the Meye and Adan labs for their support during this project. We thank Barbara Sakic and Nicky van Kronenburg for technical support. We thank Manuel Mameli, Salvatore Lecca and Stephane Ciocchi for critical review of the manuscript. This work was supported by the Dutch Research Council with NWO Veni (016.Veni.192.188) and XS (OCENW.XS21.3.076) grants to D.R.; by the European Union’s Horizon 2020 research and innovation program ERC Starting grant (804089; ReCoDE) to F.J.M.; and by the Dutch Research council (NWO) via an ENW-VIDI (VI.Vidi.203.102) grant to F.J.M.

## Authors Contributions

D.R., F.J.M., study conceptualization, project administration, supervision, manuscript writing, review and editing, funding acquisition. D.R., data acquisition, analysis and visualization. K.R data acquisition and analysis. I.G.W-D. technical assistance.

