## Supplementary Figures and Legends for "Neuropeptide Y neurons of the locus coeruleus inhibit noradrenergic system activity to reduce anxiety"

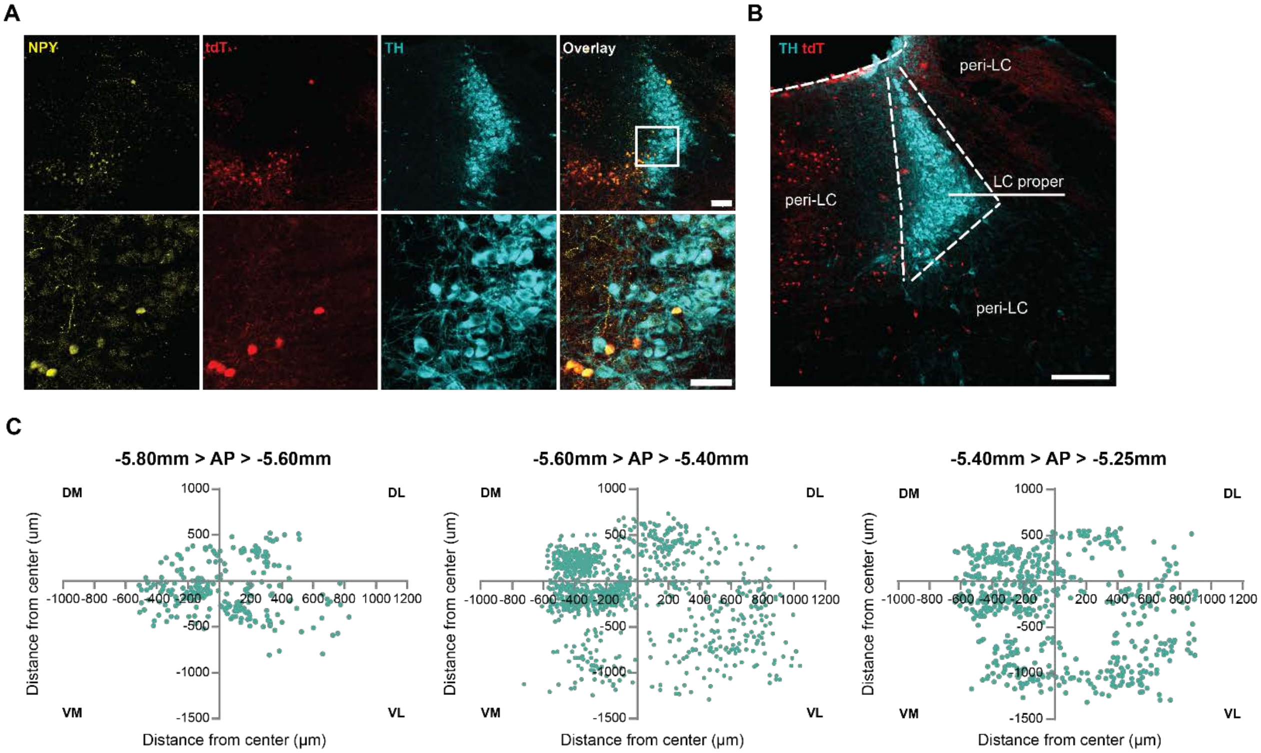


Figure S1

**Figure S1. Distribution of NPY neurons in the pericoerulean spaces and TH co-expression.**

A) Representative example of tdTomato expression (tdT, red), NPY (yellow) and $LC_{NE}$ neurons (TH, cyan) immunolabeling in coronal slices from *NPY-cre:Ai14* mice, depicting the LC. Inset: Tdt+ neurons colocalize with endogenous NPY, but not TH; Scale bar, 100µm and (inset) 50µm. B) Representative image of the LC proper, as indicated by $TH^+$ (cyan) $LC_{NE}$ cell bodies, and the pericoerulean spaces occupied by NPY-expressing ($tdT^+$, red) neurons. Scale bar, 200µm. C) The location of peri-$LC_{NPY}$ neurons, identified by tdT expression, was mapped against $LC_{NE}$ cells in the entire rostrocaudal axis containing the LC. Each data point represents a single $tdT^+$ cell, from which x (mediolateral) and y (dorsoventral) coordinates were extracted and plotted in respect to its distance (µm) from LC center. Data accumulated over three AP ranges. DM: dorsomedial, DL: dorsolateral, VM: ventromedial, VL: ventrolateral. AP, -5.80 to -5.60 mm, N=2 mice, n=239 cells; AP, -5.60 to -5.40 mm, N=7 mice, n=1004 cells; AP, -5.40 to -5.25 mm, N=6 mice, n=558 cells.

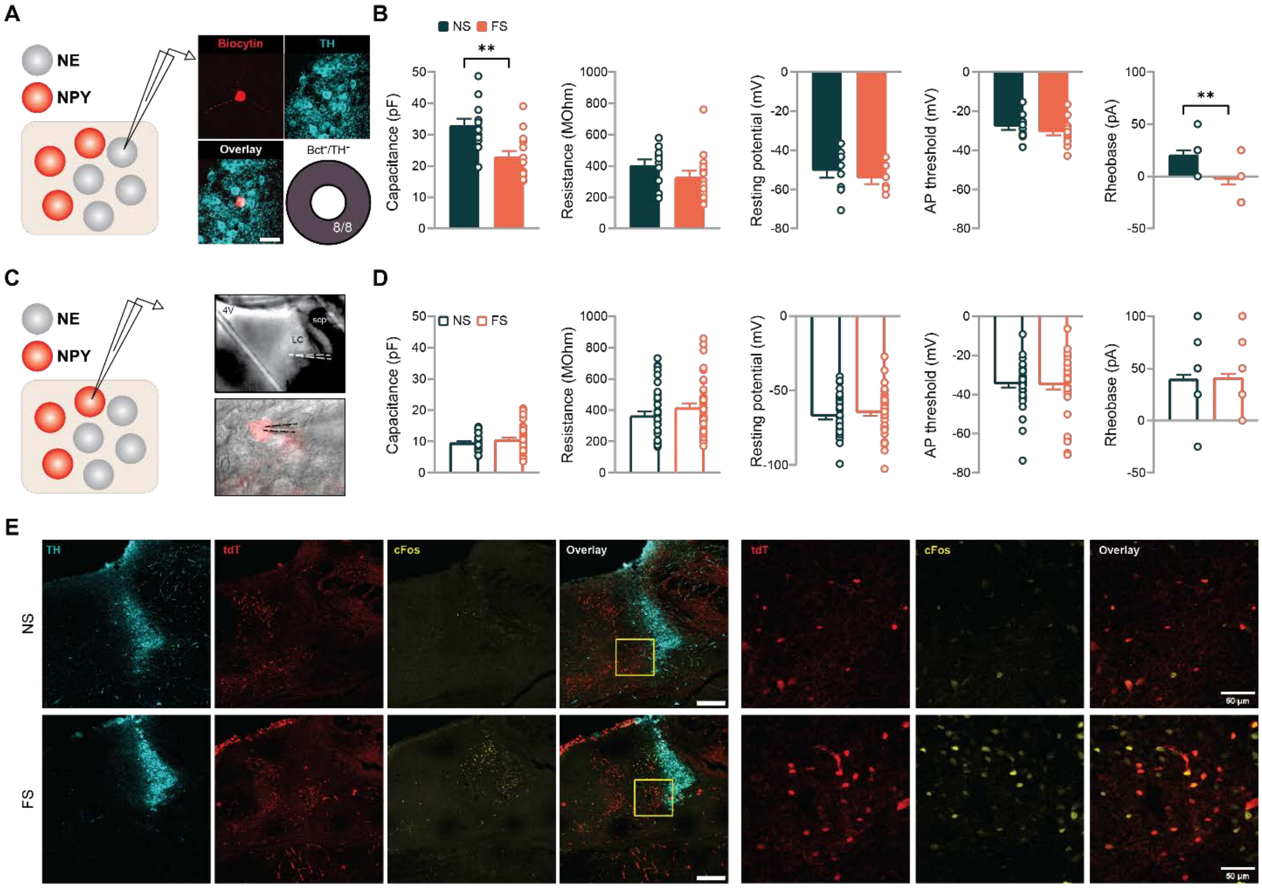


Figure S2

### Figure S2. Foot-shock stress effects on peri-LC$_{NPY}$ and LC$_{NE}$ intrinsic properties

A) Control or foot-shock-subjected *NPY-cre:Ai14* mice were sacrificed 30 min following stress exposure for electrophysiological recordings. Putative LC$_{NE}$ neurons were identified based on shape, position, and electrical properties. In a subset of recorded cells, we validated LC$_{NE}$ identity based on biocytin labeling and post-hoc TH staining. All putative LC$_{NE}$ neurons were confirmed to be TH$^{+}$ in this way (8/8). B) In LC$_{NE}$ neurons acute stress decreased cellular capacitance (NS, 32.9pF, FS, 22.9pF, unpaired t-test, t(25)=3.47, *P*=0.002) without changing resting membrane potential (NS, -50.4mV, FS, -54.3mV, unpaired t-test, t(13)=0.75, *P*=0.465), membrane resistance (NS, 404.6MOhm, FS, 328.9MOhm, unpaired t-test, t(24)=1.36, *P*=0.187) or action potential threshold (NS, -27.9mV; FS, -30.8mV, unpaired t-test, t(26)=1.28, *P*=0.210). Stress reduced rheobase in the FS group (NS, 25pA, FS, 0pA, Mann-Whitney U=41, *P*=0.006). C) Peri-LC$_{NPY}$ neurons were identified based on expression of tdTomato. D) Exposure to stress did not alter basic electrophysiological properties of peri-LC$_{NPY}$ neurons: cellular capacitance, NS, 9.6pF, FS, 10.6pF, unpaired t-test, t(84)=1.44, *P*=0.153; membrane resistance, NS, 365.9MOhm, FS, 417.2MOhm, Mann-Whitney U=682, *P*=0.144; rheobase, NS, 25pA, FS, 25pA, Mann-Whitney U=906.5, *P*=0.898; and action potential threshold, NS, -34.7mV; FS, -35.2mV, Mann-Whitney U=846, *P*=0.525. No changes in resting membrane potential were seen after stress (NS -67.4 mV; FS -64.9, unpaired t-test, t(84)=0.86, *P*=0.393). E) In a different cohort, NS and FS mice were sacrificed 90 min post-stress to examine cFos expression via immunolabeling. Representative examples of tdTomato expression (tdT, red), noradrenergic (TH, cyan) and cFos (yellow) immunolabeling in coronal slices from NS or FS mice. Inset (yellow square) marks the area shown to the left and in Fig 2F. Scale bar, 200 or 50 µm. B) NS, N=5 mice, n=12 cells, FS, N=6 mice, n=16 cells. Data depicted as mean ± SEM. * $P$ <0.05, ** $P$ <0.01. D) NS, N=10 mice, n=40 cells; FS, N=13 mice, n=46 cells. E) NS, N=4 mice, n=1572 cells FS, N=4 mice, n=867;

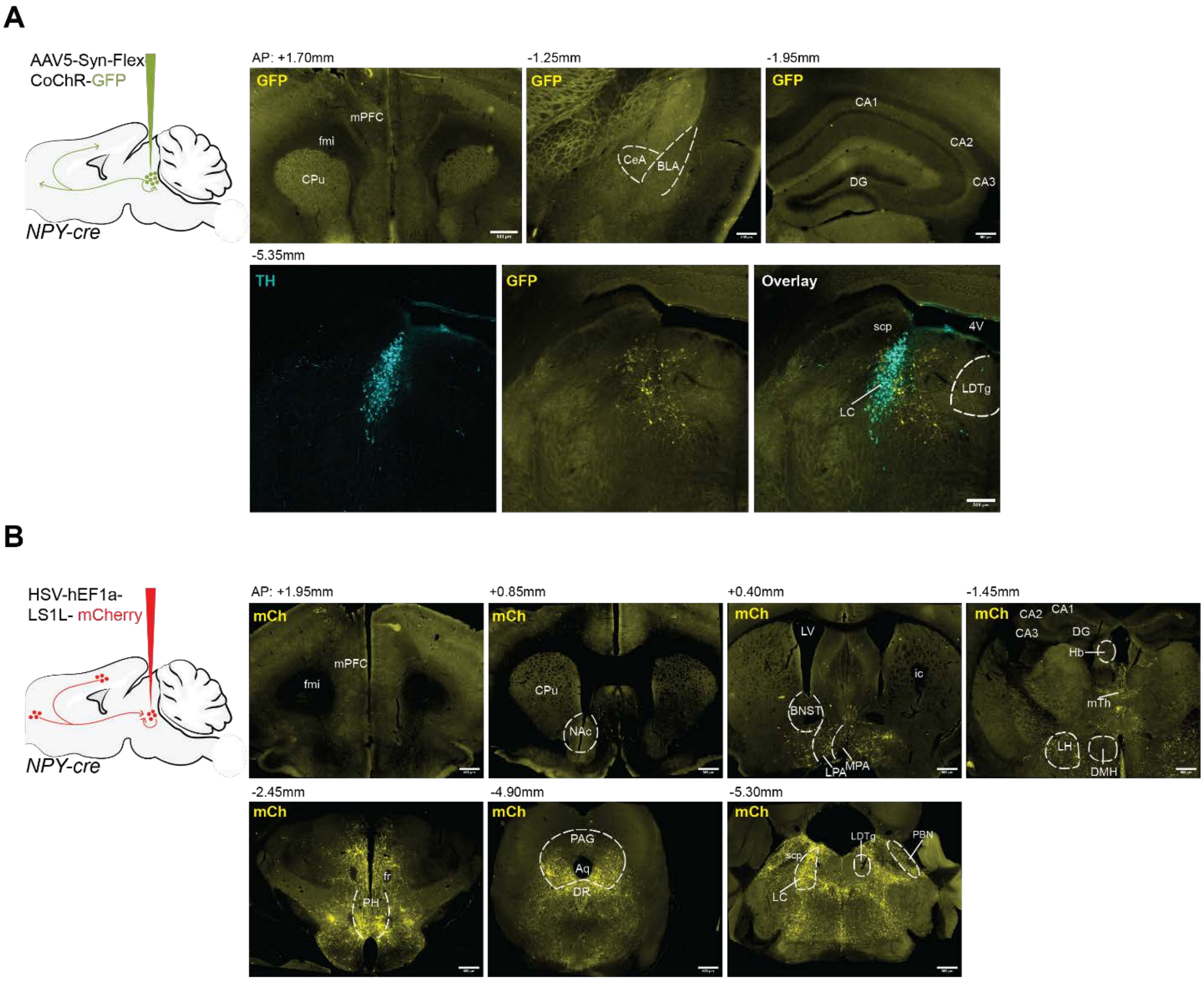


Figure S3

**Figure S3. Anterograde and retrograde tracing of peri-LC$_{NPY}$ neuroanatomical circuitry**

A) Schematic of sagittal mouse brain, depicting the location of bilateral virus injections used to drive CoChR expression in *NPY-cre* mice (N=5). Representative images show lack of NPY efferents in known LC projection fields, such as the prefrontal cortex, amygdala and the hippocampus. Conversely, GFP$^+$ cell bodies and projections are seen in the pericoerulean space (TH, tyrosine hydroxylase). B) Schematic of sagittal mouse brain, depicting the location of bilateral virus injections for retrograde labeling in *NPY-cre* mice (N=5). Several NPY afferent regions were identified (*cf.*, Table S1), with NPY neurons from hypothalamic regions and the PAG heavily innervating the LC. Aq, aqueduct; BLA, basolateral amygdala; BNST, bed nucleus of the stria terminalis; CA1, CA2, CA3 hippocampal subfields; CeA, central amygdala; CPu, caudate putamen; DMH, dorsal medial hypothalamic area; fmi, forceps minor of the corpus callosum; fr, fasciculus retroflexus; ic, internal capsule; LC, locus coeruleus; LDTg, laterodorsal tegmental nucleus; LH, lateral hypothalamic area; Hb, habenula; LPA, lateral preoptic area; LV, lateral ventricle; MPA, medial preoptic area; NAc, nucleus accumbens; PAG, periaqueductal gray; DR, dorsal raphe; PBN, parabrachial nucleus; PH, posterior hypothalamic area; mPFCL, medial prefrontal cortex; scp, superior cerebellar peduncle. Scale bar, 500μm.

| **Brain Region** | ***Cell Bodies*** |
|---|---|
| mPFC | - |
| Acb | - |
| BLA | - |
| CeA | + |
| BNST | +/- |
| HPC | - |
| LS | - |
| Hb | - |
| mTh | ++ |
| LD | + |
| PV | +/- |
| MPA | ++ |
| LPA | ++ |
| Arc | - |
| DMH | + |
| LH | ++ |
| PH | ++ |
| SN | + |
| **LC** | **++** |
| LDTg | - |
| RtTg | + |
| PBN | + |
| PAG | ++ |
| DR | ++ |

**Table S1. Retrograde labelling in *NPY-cre* mice**

Expression of mCherry$^+$ cell bodies was examined throughout the brain following bilateral injections of HSV-hEf1a-LS1L-mCherry virus targeting the LC of *NPY-cre* mice (N=5). (Pre)frontal cortical areas were devoid of retrograde labelling. Besides the pericoerulean region (*c.f.*, Fig. 1), strong expression was observed in several hypothalamic nuclei as well as in the pons. mPFC, medial prefrontal cortex; Acb, nucleus accumbens; BLA, basolateral amydgala; CeA, central amygdala; BNST, bed nucleus of the stria terminalis; HPC, hippocampus; LS, lateral septum; Hb, habenula; LD, laterodorsal thalamic nucleus; PV, paraventricular thalamic nucleus; MPA, medial preoptic area, LPA, lateral preoptic area; Arc, arcuate hypothalamic nucleus; DM, dorsal medial hypothalamus; LH, lateral hypothalamic area; PH, posterior hypothalamic area; SN, substantia nigra; LDTg, laterodorsal tegmental nucleus; LC: locus coeruleus; RtTg, reticulotegmental nucleus of the pons; PBN, parabrachial nucleus; PAG, periaqueductal gray; dorsal raphe, DR. Plus (+) and minus (-) symbols represent the degree of mCherry expression observed. Terminology for presence of retrogradely labeled NPY+ cell bodies: -no cell bodies, +/- minor presence, + moderate presence, ++ extensive presence.

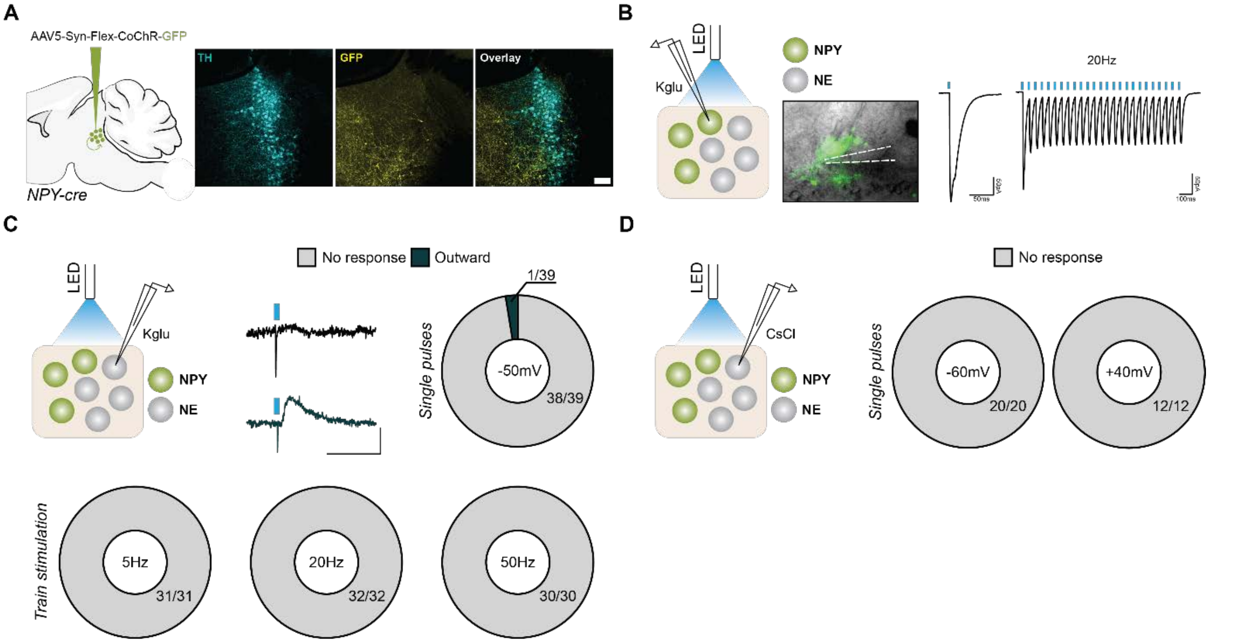


Figure S4

**Figure S4. No GABAergic or glutamatergic synaptic connectivity between peri-$LC_{NPY}$ and $LC_{NE}$ neurons**

A) Left: Schematic of sagittal mouse brain, depicting the location of bilateral virus injections used to drive CoChR expression in *NPY-cre* mice. Right: After a period of virus incubation (≥ 5 weeks) extensive CoChR innervation (GFP, yellow) was observed in the pericoerulean region (TH, cyan), in brain slices prepared for electrophysiological recordings. B) Left: Schematic representation of experimental design. In brain slices containing CoChR, we confirmed that peri-$LC_{NPY}$ neurons expressing CoChR ($GFP^+$ cell) directly respond with photocurrents to respective optical stimulation (1ms, single or 20Hz pulse train of 20 pulses), validating efficiency of the optogenetic construct. Recording pipet is indicated C) Top: In brain slices containing CoChR-expressing peri-$LC_{NPY}$ neurons, we recorded $LC_{NE}$ postsynaptic responses to 1 ms LED-mediated blue light pulses. Recordings were performed in voltage-clamp configuration, at -50mV, using a KGlu-based internal solution, to allow for detection of both glutamatergic (inward) and GABAergic (outward) responses. No postsynaptic responses to peri-$LC_{NPY}$ photostimulation were observed in the majority of $LC_{NE}$ cells (38/39), indicating next-to-null ionotropic receptor-mediated connectivity in the two populations. One cell showed a nominal outward, presumably GABA-mediated, response. Example traces are shown (scale bar: 5 mV, 10 ms). Blue rectangle: start of LED stimulation. Bottom: $LC_{NE}$ synaptic responses to pulse trains of optical stimulation paradigms were recorded. No postsynaptic responses were detected in any of the frequencies sampled (5Hz, 31/31 cells; 20Hz, 32/32; 50Hz, 30/30 cells, non-responsive). D) In brain slices containing CoChR-expressing peri-$LC_{NPY}$ neurons, we recorded $LC_{NE}$ opto-responses at -60mV or +40mV, using a CsCl-based internal solution, to increase sensitivity in detection of $GABA_AR$ (or AMPAR), or NMDA-mediated currents. No response to optostimulation in any of the recording conditions was observed. C) Single pulse: N=14 mice, n=39 cells; 5Hz: N=13 mice, n=31 cells; 20Hz: N=13 mice, n=32 cells; 50Hz: N=12 mice, n=39 cells D) -60 mV, N=6 mice, n=20 cells; +40mV, N=3 mice n=12 cells.

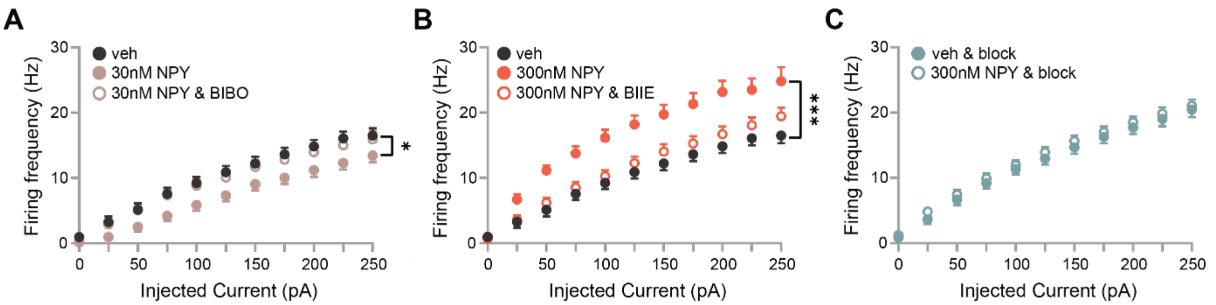


Figure S5

**Figure S5. Effects of NPY bath application on $LC_{NE}$ excitability**

A) Brain slices from wild-type C57Bl/6 mice were prepared for electrophysiological recordings. Whole-cell patch clamp recordings from $LC_{NE}$ neurons were conducted in current-clamp configuration, while 30nM NPY or vehicle were continuously perfused. In the presence of 30nM we detected fewer action potentials in response to increasing current injections as compared to vehicle, and this effect was blocked by pretreatment with the selective Y1R antagonist BIBO-3304 (1µM, Firing frequency: 2-way RM ANOVA, main treatment effect, $F(2,45)=3.31$, $P=0.045$). B) 300nM NPY led to $LC_{NE}$ hyper-excitability, which was abolished by pretreatment with the selective Y2R antagonist BIIE-0246 (1µM, Firing frequency: 2-way RM ANOVA, main treatment effect, $F(2,39)=10.3$, $P=0.0003$). C) The effect of high NPY dose was dependent on presynaptic input to $LC_{NE}$ cells, as pharmacological blockade of AMPARs and KainateRs (10µM CNQX), NMDARs (50µM D-AP5), $GABA_A$Rs (100µM picrotoxin) and $GABA_B$Rs (10µM CGP-54626) occluded NPY-driven increase in $LC_{NE}$ firing frequency (2-way RM ANOVA, main treatment effect, $F(1,32)=0.48$, $P=0.494$). A) Veh, N=4 mice, n=15 cells; 30nM, N=3 mice, n=15 cells; 30nM & BIBO-3304, N=3 mice, n=18 cells. B) Veh, N=4 mice, n=15 cells; 300nM, N=4 mice, n=11 cells; 300nM & BIIE-0246, N=3 mice, n=16 cells. C) Veh & block, N=3 mice, n=14 cells; 300nM NPY & block, N=3 mice, n=20 cells. Data depicted as mean ± SEM. * $P<0.05$. *** $P<0.001$.

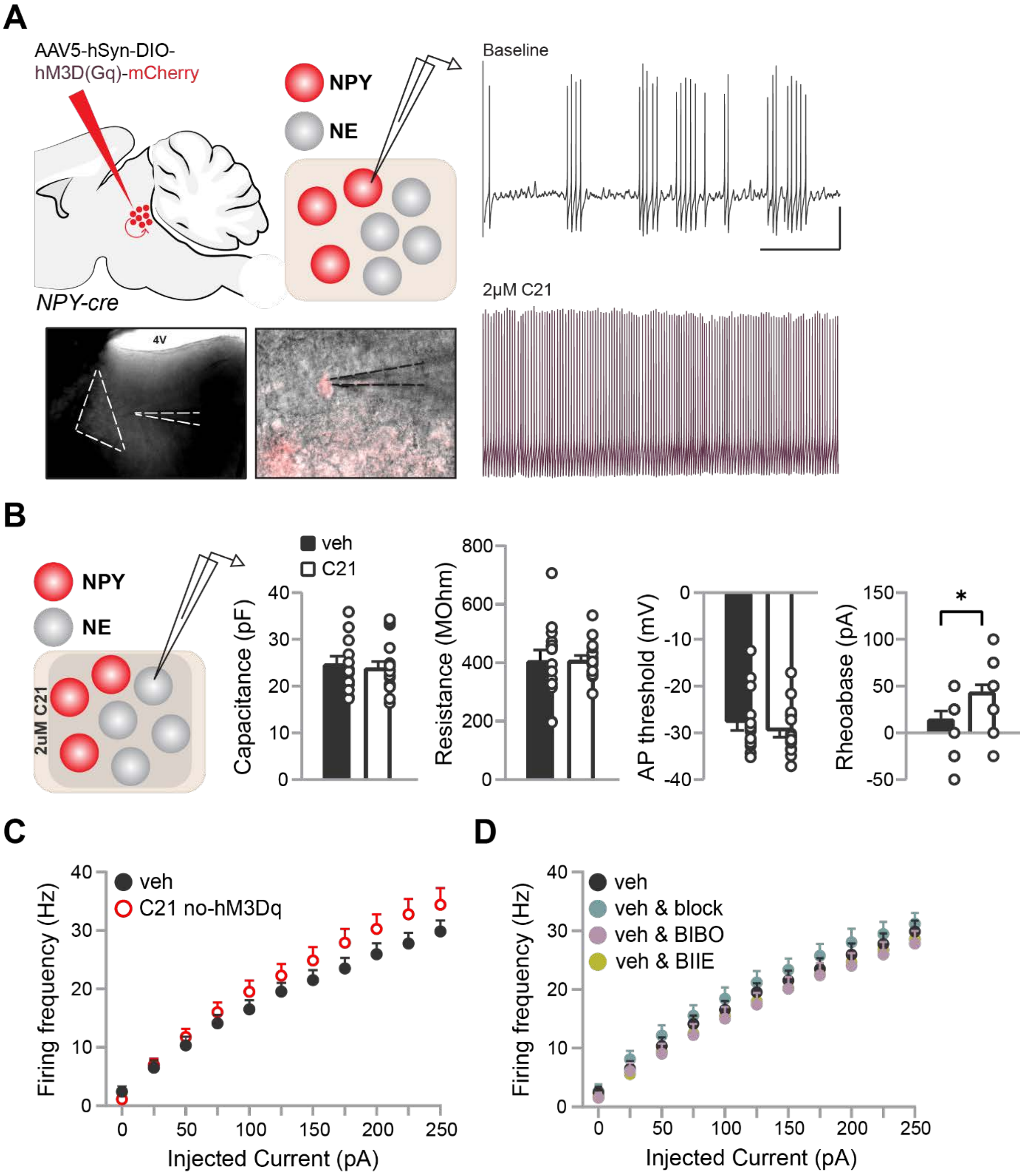


Figure S6

**Figure S6. Effects of C21 and NPY receptor antagonism on intrinsic properties and excitability of $LC_{NE}$ cells.**

A) Schematic of sagittal mouse brain depicting the location of bilateral virus injections used to drive hM3D(Gq) in *NPY-cre* mice. After a period allowing for virus expression (≥5 weeks), brain slices containing the LC were prepared and whole-cell patch clamp recordings of peri-$LC_{NPY}$ cells were conducted in current-clamp mode. Representative images of location of the recorded cell (mCherry$^{+}$) are included; LC and recording pipet are indicated (dashed lines). B) Representative traces of 2.5s of spontaneous activity of a peri-$LC_{NPY}$ cell before (top, grey) and after (bottom, magenta) bath application of the DREADD actuator C21 (2µM). Scale bar, 20 mV, 500 ms. C21 resulted in a substantial increase in the number of action potentials fired at rest. B) Brain slices from *NPY-cre* mice, bilaterally expressing the excitatory hM3Dq DREADD, were prepared for electrophysiological recordings. Whole-cell patch clamp recordings from $LC_{NE}$ neurons were conducted in current-clamp configuration, while C21 (2µM) or vehicle were continuously perfused in the bath. No effects of peri-$LC_{NPY}$ chemogenetic simulation on $LC_{NE}$ cellular capacitance (veh, 24.8pF *vs.* C21, 24.0pF, Mann-Whitney U=117.5, $P$=0.716), membrane resistance (veh, 408.1MOhm *vs.* C21, 407.9MOhm, unpaired t-test, t(29)=0.04, $P$=0.965), or action potential threshold (veh, -27.8mV *vs.* C21, -29.6mV, Mann-Whitney U=108.5, $P$=0.484) were observed. In support of reduced $LC_{NE}$ excitability (*cf.*, Fig. 3F), C21 increased the current necessary for $LC_{NE}$ cells to exceed their action potential threshold and fire (Rheobase: veh,15.0pA *vs.* C21, 44.1pA, Mann-Whitney U=65, $P$=0.013). B) In brain slices of *NPY-cre* mice that did not express hM3Dq, we recorded $LC_{NE}$ firing frequency in response to increasing currents in presence of C21. C21 alone did not alter $LC_{NE}$ excitability (2-way RM ANOVA, main effect of treatment, F(1,23)=1.38, $P$=0.252), controlling for effects of unspecific binding of the DREADD actuator. C) Effects of receptor antagonism on $LC_{NE}$ firing patterns. In vehicle-treated slices, pharmacological blockade of AMPAR-, NMDAR-, $GABA_AR$ and $GABA_BR$-mediated input or antagonism of Y1R and Y2R had no effect on $LC_{NE}$ firing (2-way RM ANOVA, main effect of treatment, F(3,49)=0.71, $P$=0.550). B) Veh, N=4 mice, n=15 cells; C21, N=4 mice, n=17 cells. C) Veh, N=4 mice, n=15 cells; C21, N=2 mice, n=10 cells. D) Veh, N=4 mice, n=15 cells; Veh & block, N=2 mice, n=14 cells; Veh & BIBO-3304, N=3 mice, n=13 cells; Veh & BIIE-0246, N=2 mice, n=11 cells. Data depicted as mean ± SEM. * $P$ <0.05.

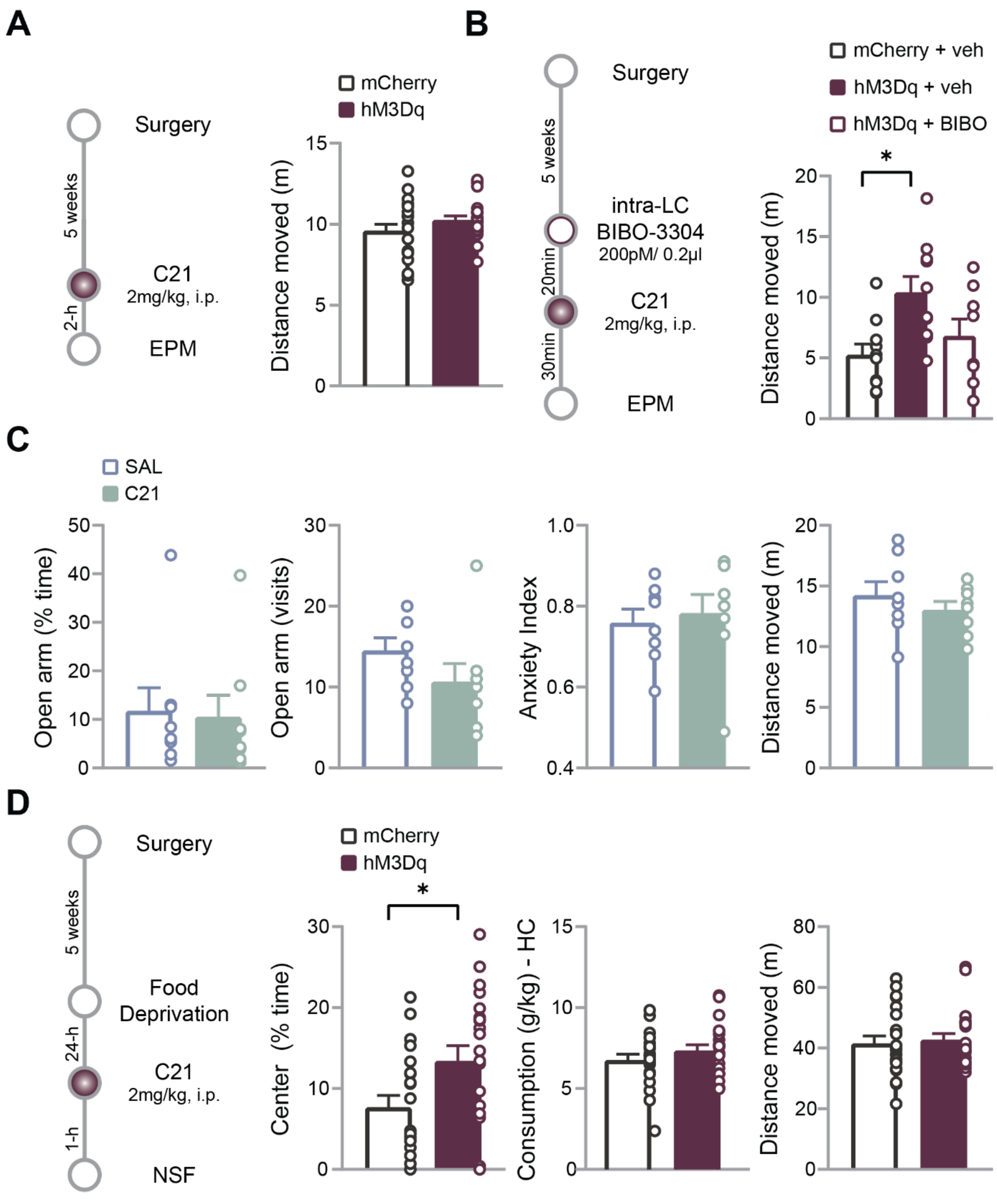


Figure S7

**Figure S7. Behavioral outcomes of peri-LC$_{NPY}$ chemogenetic activation and C21 administration**

A) Experimental design for *in vivo* chemogenetic manipulations: After 5 weeks allowing for virus expression, hM3Dq and mCherry groups were systemically administered C21 (2mg/kg, i.p.) and 2-h later subjected to the elevated plus maze (EPM) task. Chemogenetic activation of peri-$LC_{NPY}$ neurons had no consequences on general locomotor activity, as measured by total distance moved at the EPM task (mCherry, 9.6m; hM3Dq, 10.3m; unpaired t-test, t(40)=1.37, *P*=0.178). B) However, in a subsequent EPM experiment, in animals that received intra-LC micro-infusions via cannula, C21 (2mg/kg) altered locomotion in hM3Dq & veh group, which displayed significantly increased distance moved *vs.* mCherry & veh controls (*P*=0.012). No other statistically significant group differences were detected (mCherry & veh *vs.* hM3Dq & BIBO, *P*=0.653; hM3Dq & veh *vs.* hM3Dq & BIBO, *P*=0.119). The observed discrepancy between A) and B) could be due to differences in experimental design, and in particular the time interval between C21 administration and EPM (2h *vs.* 30min, respectively). C) Controlling for non-specific, off-target effects, administration of C21 (2mg/kg, i.p.), did not affect EPM performance nor general locomotion as compared to vehicle (saline), in a separate cohort of wild-type C57BL/6 mice that did not express the hM3Dq-carrying viral construct: time spent in open arms (% of total exploration time): SAL, 11.7%; C21, 10.5%; Mann-Whitney, U=27, *P*=0.645); open arms entries: SAL, 14.5; C21, 10.6; unpaired t-test, t(14)=1.39, *P*=0.186); anxiety index: SAL, 0.76; C21, 0.78; unpaired t-test, t(14)= 0.41, *P*=0.688); distance moved: SAL, 14.2m; C21, 13.0m; unpaired t-test, t(14)=0.90, *P*=0.384). D) After 5 weeks allowing for hm3Dq expression, mice were food deprived (24-h) and systemically administered the DREADD agonist compound 21 (C21, 2mg/kg, i.p.). One hour after after C21 injection, mice we subjected to the novelty suppressed feeding (NSF) task. hM3Dq group spent more time at the center of the NSF arena, containing a familiar food source, vs. mCherry controls (Duration Center: mCherry, 7.7s; hM3Dq, 13.4s; Mann-Whitney U=137, *P*=0.035). Chemogenetic activation of peri-$LC_{NPY}$ neurons did not affect general consummatory behavior, as no group differences in home-cage food intake were seen (mCherry, 6.76 g/kg; hM3Dq, 7.35 g/kg; unpaired t-test, t(40)=1.14, *P*=0.261). No unspecific effects of C21 in general locomotion were seen (Distance moved: mCherry 41.5m, hM3Dq, 42.7m; Mann-Whitney U=202, *P*=0.654). A, D) mCherry, N=21; hM3Dq, N=21. B) mCherry & veh, N=10; hM3Dq & veh, N=10; hM3Dq & BIBO-3304, N=8. C) SAL, N=8; C21, N=8. Data depicted as mean ± SEM. * *P* <0.05.

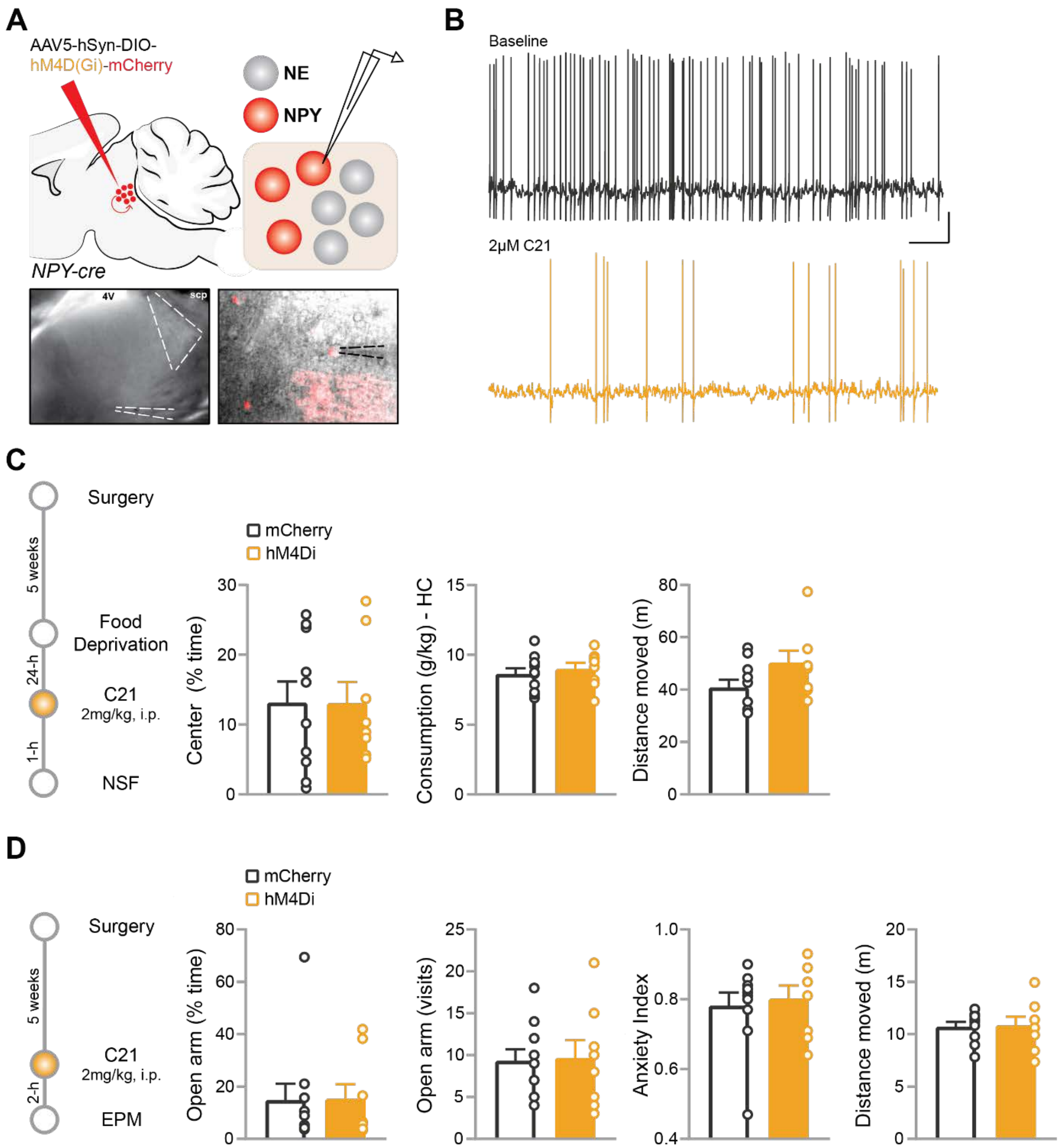


Figure S8

**Figure S8. Behavioral outcomes of peri-LC$_{NPY}$ chemogenetic inhibition**

A) Schematic of sagittal mouse brain depicting the location of bilateral virus injections used to drive hM4D(Gi) in *NPY-cre* mice. After a period allowing for virus expression (≥5 weeks), brain slices containing the LC were prepared and whole-cell patch clamp recordings of peri-LC$_{NPY}$ cells were conducted in current-clamp mode. Representative images of location of the recorded cell (mCherry$^{+}$) are included; LC and recording pipet are indicated (dashed lines). B) Representative traces of 30s of spontaneous activity of a peri-LC$_{NPY}$ cell before (top, grey) and after (bottom, orange) C21 (2μM) bath application (Scale bar, 20mV, 2.5 s). C21 resulted in a substantial decrease in the number of action potentials fired at rest. C) Experimental design for *in vivo* chemogenetic manipulations: after 5 weeks allowing for virus expression, mice were food deprived (24-h) and systemically administered the DREADD agonist compound 21 (C21, 2mg/kg, i.p.). One hour after C21 injection, mice we subjected to the novelty suppressed feeding (NSF) task. In hM4Di mice, C21 did not alter time spent at the center of the NSF arena as compared to mCherry controls (Duration Center: mCherry, 13.1s; hM4Di, 13.0s; unpaired t-test t(16)=0.01, *P*=0.988). C21 effects on consummatory behavior were specific for the anxiogenic context (*cf.*, Fig. 4H), as chemogenetic inhibition of peri-$LC_{NPY}$ neurons did not affect general consummatory behavior (Home-cage food intake: mCherry, 8.62 g/kg; hM4Di, 8.97 g/kg; unpaired t-test, t(16)=0.57, *P*=0.575). Finally, C21 did not affect general locomotion (Distance moved: mCherry, 40.8m; hM4Di, 50.3m; unpaired t-test, t(16)=1.78, *P*=0.094). D) hM3Di and mCherry groups were systemically administered C21 (2mg/kg, i.p.) and 2-h later subjected to the elevated plus maze (EPM) task. In contrary to NSF, we observed no anxiogenic effects of peri-$LC_{NPY}$ silencing on EPM performance: time spent in open arms (% of total exploration time): mCherry, 14.8%; hM4Di, 15.3%; Mann-Whitney U=38, *P*=0.897); open arms entries: mCherry, 9.3; hM4Di, 9.6; unpaired t-test, t(16)=0.13, *P*=0.897); anxiety index: mCherry, 0.78; hM4Di, 0.80; unpaired t-test, t(16=0.37, *P*=0.716). C21 did not affect general locomotor activity in EPM (Distance moved: mCherry, 10.7m; hM4Di, 10.8m; unpaired t-test, t(16)=0.12, *P*=0.906). C, D) mCherry, N=10; hM4Di, N=8. Data depicted as mean ± SEM
